# Sensorimotor Theta Oscillations Coordinate Speech Movements

**DOI:** 10.1101/2025.10.09.681482

**Authors:** Yitzhak Norman, Loren M. Frank, Edward F. Chang

**Author notes:** Corresponding author. EFC.

## Abstract

Fluent speech depends on precisely timed motor commands that coordinate rapid transitions between successive articulatory gestures. Using direct cortical recordings, we identified a prominent sensorimotor theta oscillation (6–10 Hz) that supports this coordination. During articulation, premotor speech circuits exhibited enhanced theta phase coherence, with elevated population activity near theta troughs. The oscillation’s frequency remained remarkably stable across varying speech rates and cognitive states, consistent with an intrinsically generated rhythm. Vocal-tract kinematics revealed pulse-like movements at 6–10 Hz, tightly coupled to cortical theta phase. At a mesoscopic scale, theta cycles structured sequential sensorimotor activations encoding articulatory gestures, with syllable identity optimally decodable following theta troughs. These findings identify theta oscillations as an intrinsic timing mechanism that coordinates the distributed and synergistic motor control underlying fluent speech.

## Main Text

Speech is a defining human behavior, enabling us to express an unlimited range of thoughts by flexibly combining a limited set of basic phonetic elements. To speak fluently, the brain must generate rapid and precisely timed motor commands—at the scale of tens of milliseconds (*1*)— that coordinate the activity of nearly 100 muscles within the vocal tract (*2*). Each speech sound is produced through coordinated, synergistic movements of the tongue, lips, jaw, and other articulators, which dynamically shape specific constriction gestures (*3*). In fluent speech, transitions between these gestures are swift and seamless.

While previous research has shown that distributed neural activations within the ventral sensorimotor cortex (vSMC) play a central role in driving articulatory gestures (*1*, *4–10*), the mechanism that ensures precise temporal coordination among the underlying neuronal populations remains enigmatic. Specifically, how does the brain endogenously activate these distributed neuronal representations in unison at precisely the right moment? How does it orchestrate successive motor commands to enable smooth transitions from one constriction gesture to the next? Uncovering this coordination mechanism is essential not only for deepening our understanding of motor control circuits underlying speech and other dexterous behaviors in the human brain, but also for advancing novel therapeutic strategies for speech and motor disorders and improving the precision of brain-computer interface technologies (*11*) .

Insights from animal models point to a potential coordination mechanism rooted in intrinsic neural oscillations. In primates and rodents, rhythmic activity—particularly in the 6-10 Hz theta/alpha range—has been shown to synchronize neuronal spiking and population-level processing within and across brain regions (*12–19*). Across species, such oscillations have been implicated in the temporal coordination of diverse motor behaviors (*20–24*), including non-human primate vocalization (*25*) and human reading saccades (*26*). In the rodent hippocampus, theta oscillations organize information flow (*27–29*) and align hippocampal activity with locomotor (*30*), olfactory (*31*, *32*), somatosensory (*33*, *34*) and prefrontal representations on timescales of ∼20-50 milliseconds (*13*, *15*, *35*). Together, these findings suggest that intrinsic oscillatory dynamics within the theta range can facilitate temporally precise interactions among spatially dispersed neuronal populations.

In humans, converging evidence links theta activity to large-scale interregional communication (*36*, *37*), most notably during spatial navigation (*38–42*) and medial temporal lobe memory processing (*43–45*). Yet whether theta oscillations organize the processing of specific information streams remains unclear. Here, we tested whether theta rhythmicity supports the coordination of distributed motor control in the human brain—specifically, the motor control of speech, which relies on extensive premotor networks to orchestrate the rapid and complex muscle synergies underlying articulatory gestures (*5*, *10*, *46*).

Using high-density electrocorticographic (ECoG) recordings in neurosurgical patients, we identify a robust 6-10 Hz theta oscillation in the local field potentials (LFPs) of the vSMC and adjacent regions. This intrinsic neural rhythm exhibits stable power and frequency during ongoing articulation and shows coherent phase across widely distributed, speech-responsive sites. Crucially, during fluent speech, both vocal tract kinematics and cortical representations of articulatory movements are tightly coupled to specific phases of the oscillation. Taken together, our findings suggest that this circuit-level theta rhythm functions as an intrinsic timing mechanism—a shared rhythmic scaffold—that synchronizes premotor neuronal populations to coordinate the rapid and precisely timed articulatory movements required for fluent speech.

## Results

High-density electrocorticographic (ECoG) recordings were obtained from 17 patients undergoing intracranial epilepsy monitoring (5 females, 12 males, age range: 22-60, M=36.3). Participants spoke hundreds of sentences from the MOCHA-TIMIT corpus (*47*), covering the full range of the English phonetic inventory. Sentences were displayed on a screen, and after receiving a go cue, participants read them aloud at their natural speaking rate (Fig. 1A and Methods for task details). In addition, participants completed control tasks that included spontaneous speech in natural conversation, passive listening to sentences, and silent rest.

**Fig. 1.**
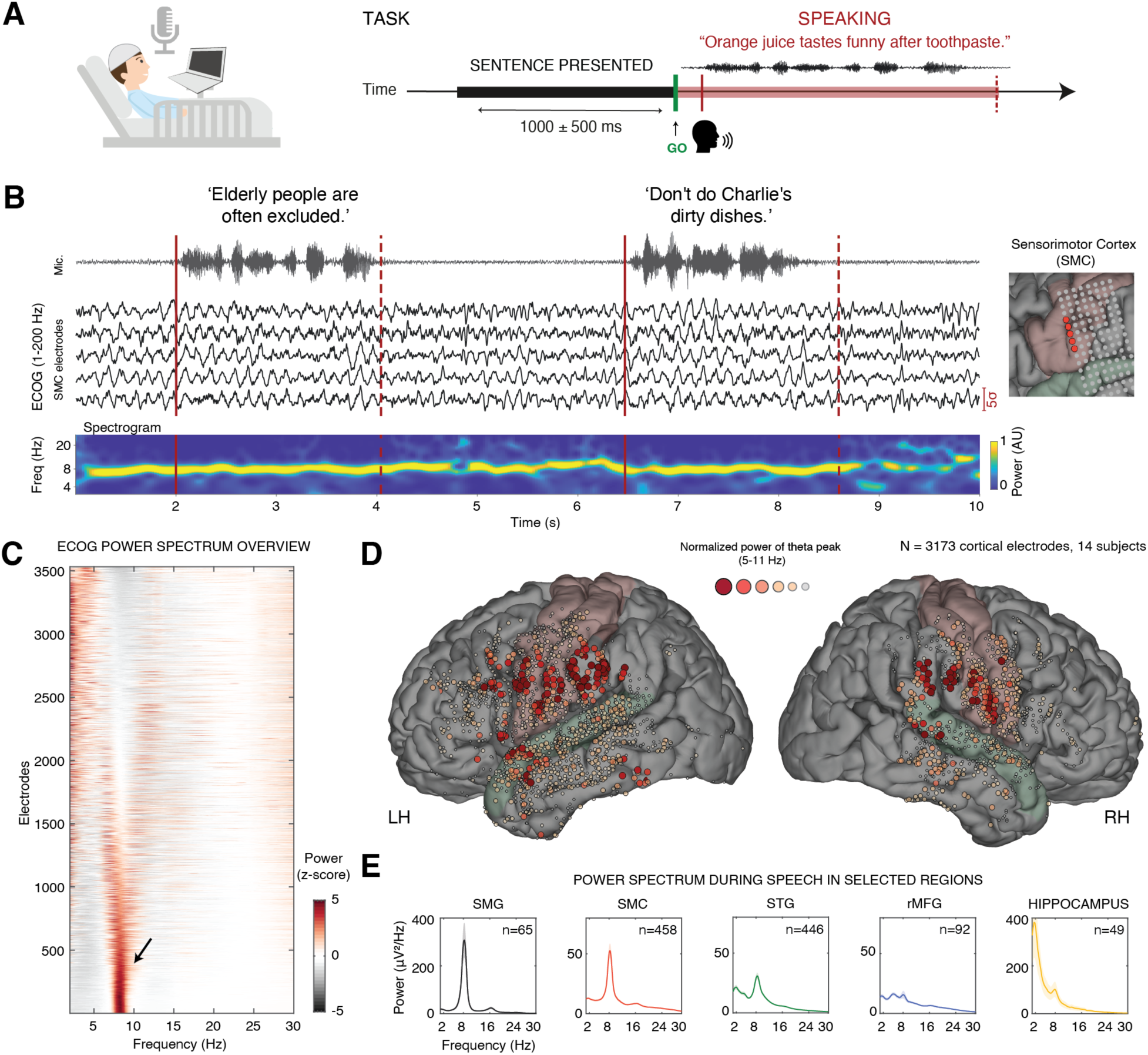
Prominent 8 Hz neural oscillation centered in the sensorimotor cortex during speech production. **(A)** Intracranial electrodes were implanted in the cortex and hippocampus as part of neurosurgical treatment for medically intractable epilepsy. During the task, participants spoke hundreds of English sentences covering the full phonetic inventory. **(B)** Example of 8 Hz theta oscillation during speech production, as it appears in five speech-responsive sensorimotor sites in one representative patient (wideband ECOG signal, filtered between 1-200Hz; right panel shows electrode location). The spectrogram at the bottom shows peak frequency around 8 Hz. This ongoing rhythm was detectable also during inter-trial silence period, pointing to its ongoing, intrinsic nature. Voice amplitude is shown at the top. **(C)** An overview of the power spectral density during speech production across 3,599 recording sites in 14 patients. A prominent power concentration in the high theta range (6–10 Hz) is evident across a major portion of the electrodes, with 41% of them displaying a single, distinct, theta peak (black arrow). **(D)** Cortical electrodes were color-coded according to the magnitude of their theta-band spectral peak. An evident cluster is observable over the sensorimotor cortex (SMC; painted in light pink) and surrounding areas, including the supramarginal gyrus (SMG), inferior frontal gyrus (IFG) and superior temporal gyrus (STG; painted in pale green). **(E)** Mean power spectrum during speech across selected regions of interest. From left to right: speech-responsive SMG sites (black, n=65), speech-responsive SMC sites (red, n=458), speech-responsive STG sites (green, n=446), speech non-responsive sites in rostral MFG (blue, n=92), and hippocampal electrodes (yellow, n=49). Notably, a distinct and robust 8 Hz spectral peak is observed in the SMC, SMG, and STG—cortical regions involved in speech motor control. A similar spectral peak is also evident in the hippocampus, indicating spatial propagation of this oscillation into subcortical structures. As a control region, non-responsive electrodes in the rostral MFG do not exhibit this prominent 8 Hz peak. Shaded areas represent least squares mean estimates ± SE across electrodes, derived from a linear mixed-effects model.

A total of 3,599 electrodes were analyzed, of which 1,438 were identified as speech-responsive sites—defined as sites exhibiting a significant increase in high-gamma (HG) amplitude during reading aloud compared to either pre- or post-utterance silent periods (P < 0.05, FDR corrected, see Fig. S1 and Methods). During articulation, we observed a prominent LFP oscillation in speech-responsive sensorimotor areas and adjacent regions. This oscillation was robust and widespread, persisted throughout the articulatory process, and could often be detected during inter-trial silence periods—consistent with an ongoing, intrinsically generated rhythm (Fig. 1B).

To characterize the spectral signature of this oscillation across recording sites, we computed the power spectral density for each electrode within a time window of –1 to 5 sec relative to speech onset (see Methods). Fig. 1C summarizes the resulting spectra, focusing on frequencies below 30 Hz. We observed a distinct theta-band peak centered around 8 Hz (black arrow), rising above the 1/f background, was observed in 41% of all cortical electrodes—defined by a peak prominence > 3σ relative to the z-scored, 1/f-corrected power spectrum (see Methods). Additional electrodes exhibited a broader, yet still pronounced, power concentration within the high-theta/lower-alpha range (6–10 Hz). Fig. 1D shows the anatomical locations of these electrodes, color-coded by the magnitude of the detected spectral peak. The oscillation was particularly prominent in the vSMC (see Fig. S1 for example electrodes) and neighboring regions.

To assess whether the spatial distribution of this rhythm aligns with speech production circuits, we computed the mean power spectrum across speech-responsive electrodes in key regions of interest (Fig. 1E). This analysis revealed a narrow-band spectral peak centered around 8 Hz during articulation, consistently observed across multiple nodes of the speech production network, including the vSMC, supramarginal gyrus (SMG), superior temporal gyrus (STG), and more distal structures such as the hippocampus. To further assess functional specificity, we compared theta power during speech between vSMC electrodes and non-responsive electrodes in a nearby cortical region outside the speech production network—specifically, the rostral middle frontal gyrus (rMFG). This within-subject comparison revealed significantly stronger theta power in vSMC (6– 10 Hz power, vSMC vs. rMFG: t(548) = 2.63, P = 0.009; mixed-effects analysis, see Methods).

We hypothesize that the prominent vSMC-centered oscillation observed during speech serves as an intrinsic timing mechanism—a circuit-level “metronome”—that provides a rhythmic scaffold for synchronizing distributed premotor populations controlling speech movements. This synchronization is essential for producing the precisely timed vocal-tract gestures that underlie fluent and continuous speech.

This hypothesis leads to several key predictions, which we examine in the following sections: (1) *Frequency stability:* to provide a reliable rhythmic reference, the oscillation should maintain a stable frequency during articulation. (2) *Phase coherence:* to coordinate a shared central rhythm, theta phase should be coherent across spatially dispersed speech-production sites. (3) *Activity modulation:* local population firing should exhibit significant modulation by theta phase. (4) *Behavioral correlation:* vocal tract kinematics are expected to show transient movements phase-locked to theta. (5) *Mesoscopic-level orchestration*: patterns of distributed premotor activations, encoding specific articulatory gestures, should be spatiotemporally organized by the theta cycle.

### Intrinsic stability of the sensorimotor theta rhythm

The first prediction of our intrinsic scaffold hypothesis is that the oscillation’s central frequency should remain stable despite moment-to-moment fluctuations in speech syllable rate—providing a consistent temporal reference that can structure the behavior rather than being passively driven by it.

To test this, we pooled fluently articulated sentences within each subject (excluding those with speech errors or repairs) and grouped utterances into slow/medium/fast tertiles based on syllable rate (mean ± SD: slow, 2.9±0.5; medium, 3.8±0.5; fast, 4.7±0.6 syl/s). Power spectra from speech-responsive SMC sites were computed separately for each tertile (Fig. 2A). We then applied parametric spectral decomposition using the FOOOF algorithm (*48*) to separate the aperiodic 1/f component from the oscillatory components (see Methods). Across rates, a robust peak was consistently present around 8 Hz (Fig. 2B). The central frequency of peaks within the broader theta range (5-11Hz) did not vary with speech rate (mixed effects: F(2,1265) = 1.66, P = 0.19; Fig. 2C). This stability, despite substantial variation in syllable rate, supports the interpretation that this oscillation reflects an intrinsic, self-organized rhythm rather than a passive byproduct of articulation or its acoustic consequences; were it a passive byproduct, the frequency would be expected to track speaking rate.

**Fig. 2.**
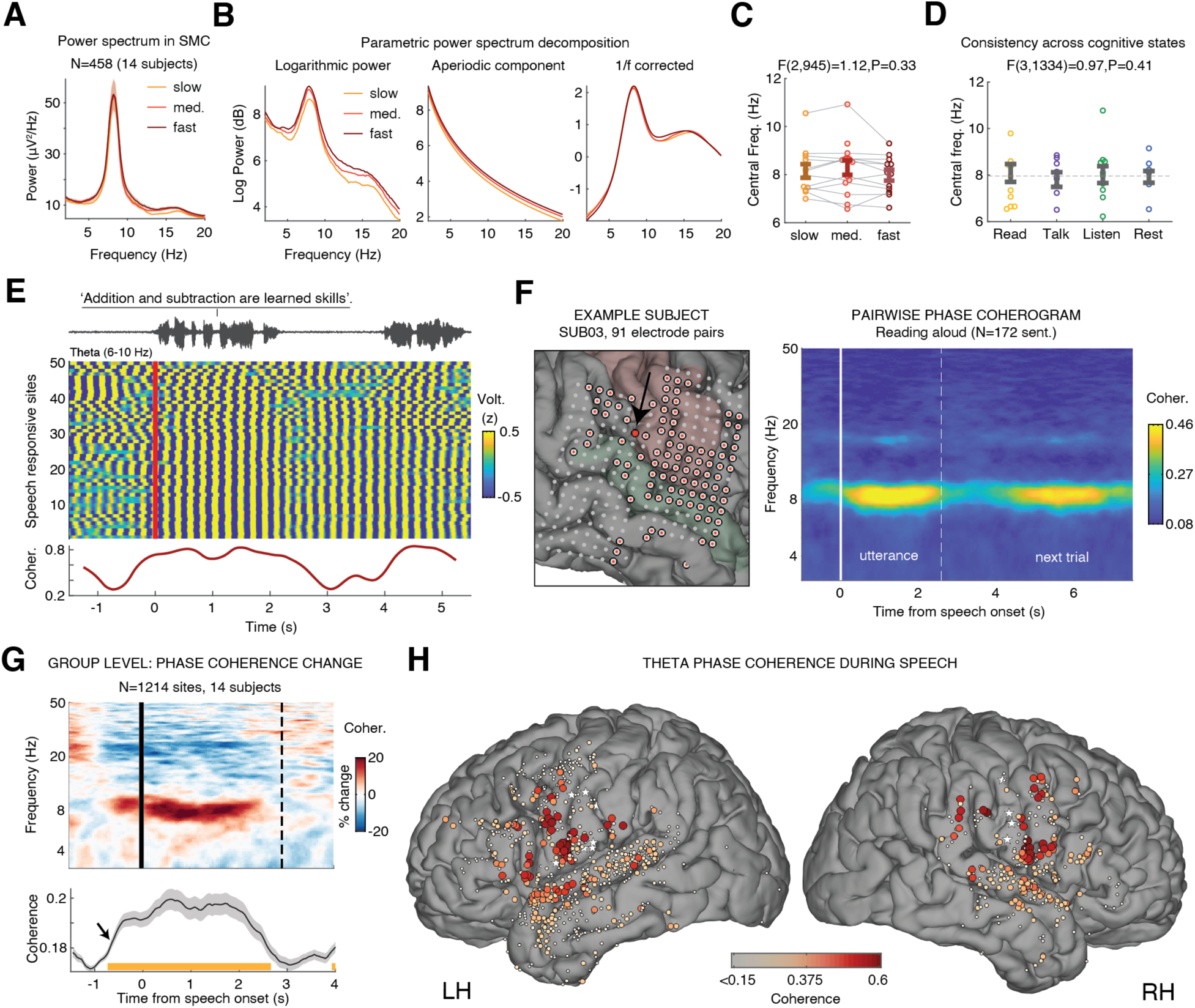
Intrinsic oscillatory stability and inter-regional theta phase coherence during speech. **(A)** Power spectrum during slow (bottom third), medium (middle third), and fast (top third) utterances. Despite substantial variation in syllable rate (2.9, 3.8, and 4.7 syl/s, respectively), the power spectrum remained markedly stable, peaking consistently around 8 Hz. **(B)** Parametric decomposition of the power spectrum in SMC sites (*53*) into aperiodic (1/f) and oscillatory components revealed a clear oscillatory peak in the high theta range (6-10 Hz). **(C)** The central frequency of theta-range peaks did not differ significantly across speech rates (F(2,890)=1.61, P=0.20, mixed-effects analysis; grand average: 8.2 ± 0.16 Hz). (D) The oscillation’s central frequency likewise remained stable across behavioral states—reading aloud, spontaneous speech, passive listening, and silent rest (F(3,1244)=0.36, P=0.78, mixed-effects analysis). **(E)** Example from a representative patient showing increased inter-site theta phase coherence during articulation. Top: voice amplitude. Bottom: sliding-window phase coherence (0.5-s windows, 0.05-s step) across 50 speech-responsive sites. **(F)** Averaged pairwise coherograms between a selected vSMC site exhibiting a prominent theta peak (black arrow) and all other speech-responsive sites in a representative patient (91 electrode pairs, marked with red dots; N = 172 sentences). A selective increase in phase coherence is evident in the 6–10 Hz range during articulation (dashed vertical line marks the median utterance duration). **(G)** Top: Group-level analysis showing the grand average change in phase coherence during articulation, computed across all speech-responsive sites relative to a selected vSMC site in each patient that exhibited a prominent theta oscillation. Bottom: phase coherence in the 6–10 Hz range increased significantly relative to the pre-speech silent period [-1 to 0] (timepoint-by-timepoint mixed-effects analysis, P < 0.05, FDR-corrected). Note the preparatory increase in coherence preceding speech onset (black arrow). Shaded area indicates the SE of the mixed-effects model estimates. **(H)** Anatomical distribution of theta phase coherence among speech-responsive electrodes during articulation (1,214 pairs, 14 subjects). White stars mark the selected theta sites used as reference for the pairwise coherence computation.

A silent-miming control in five subjects further ruled out acoustic feedback as the source of the oscillation: the oscillation was evident during both speaking aloud and silent miming (articulatory movements performed without sound), with no significant changes in central frequency, peak width or height (Fig. S2A-E).

Participants also completed control tasks, including spontaneous speech production (autobiographical storytelling), passive listening, and silent rest (see Methods). Applying the FOOOF algorithm to power spectra from SMC/SMG sites during these tasks showed that the central frequency of the sensorimotor oscillation did not differ significantly across behavioral states, further supporting its intrinsic origin (Fig. 2D; see Fig. S3 for more detail). Notably, relative to both passive listening and silent rest recorded outside the task, during articulation the oscillation showed a significantly narrower spectral peak—reflecting reduced frequency variability (mixed effects analysis: speaking-vs-listening: F(1,558)=4.51, P = 0.02; speaking-vs-silence: F(1,406)=7.68, P = 0.006; Fig. S3). This suggests that the rhythm becomes more periodic during articulation and relaxes during motor idling states.

Previous studies that investigated sensorimotor oscillations during other (more discrete) motor behaviors have reported mixed patterns of event-related synchronization and desynchronization (aka, “mu suppression”) time-locked to movement onset (*49–52*). To determine whether the oscillation observed here showed a more transient event-related power modulation relative to speech onset, we computed peri-utterance wavelet spectrograms in SMC and SMG electrodes (Fig. S4; see Methods). We analyzed the spectrograms using two complementary normalization techniques: relative power, in which power at each frequency is normalized by the total power across all frequencies—highlighting the relative shape of the spectrum at each time point; and baseline normalization, showing power changes relative to the immediate pre-speech silent interval.

Both analyses underscored the intrinsic stability of the oscillation. Relative power revealed sustained and focal theta-band activity throughout articulation, whereas baseline-normalized power showed minimal speech-related modulation—a stability that stands out against the pronounced power reductions observed in neighboring mu/beta frequencies. Quantifying the temporal profile of power modulation within a narrow band around the SMC’s central theta frequency (8.2 ± 1 Hz) revealed no significant change during articulation (min P = 0.15; timepoint-by-timepoint mixed-effects analysis vs. pre-speech baseline [–1 to 0], FDR-corrected). In contrast, adjacent frequencies—particularly in the higher mu range (11–13 Hz)—exhibited robust and significant power reductions during articulation (P < 0.05, FDR-corrected; Fig. S4).

Together, these spectral analyses point to an intrinsically-generated theta rhythm that remains stable across diverse speech behaviors and becomes more spectrally focused during articulation, making it well suited to serve as a reliable rhythmic coordinator for continuous speech.

### Speech-triggered increase in theta phase coherence

Having confirmed power and frequency stability, we next examined whether the *relative phases* across sites were also consistent with this oscillation functioning as a shared temporal scaffold for premotor neuronal coordination. We filtered the raw ECoG signal within the theta range (6–10 Hz) and time-locked it to speech onset. Figure 2E shows the theta-band voltage traces from 50 speech-responsive electrodes recorded simultaneously during one sentence in a single subject. During articulation, the ongoing voltage fluctuations exhibited rhythmic alignment across sites— indicative of large-scale, circuit-wide phase synchronization.

To quantify this phenomenon, we computed inter-electrode phase coherence (i.e., peri-speech coherograms) across 3–50 Hz frequencies, separately for each subject. The analysis was performed pairwise, comparing each speech-responsive electrode to a reference vSMC electrode exhibiting a prominent theta peak (Fig. 2F; see Methods). To minimize volume conduction effects, neighboring channels (<15 mm apart) were excluded from the analysis (Fig. S5).

This revealed a significant increase in phase coherence during speech, specific to the theta band (P<0.05, timepoint-by-timepoint mixed effects analysis relative to the pre-speech baseline, N=1214 electrode-pairs across 14 subjects, FDR-corrected; Fig. 2G). The spatial extent of this theta-band phase coherence was broad, encompassing multiple sites within the speech production network, with strongest coherence localized to the bilateral vSMC and extending into prefrontal, supramarginal, and superior temporal regions (Fig. 2H).

Notably, this mesoscopic, speech-related increase in phase coherence was significantly stronger during production than during passive listening—arguing against an acoustic origin. Moreover, comparable coherence magnitude was observed during spontaneous storytelling, indicating that the effect generalizes to natural, self-generated speech beyond reading (Fig. S5D-E).

While coherence quantifies the strength of coupling, it does not distinguish whether sites oscillate in synchrony or with systematic phase offsets. To disambiguate whether the observed coherence reflected true synchrony (zero phase) or consistent phase offsets across sites, we quantified the mean phase difference of each speech-responsive electrode during articulation relative to the most ventral SMC site in each subject. Across all speech-responsive sites, we observed a bimodal distribution of phase differences: the majority exhibited near-zero phase coupling (0° ± 45°; 827/1328 electrodes, 62%), while a substantial subset showed antiphase coupling (180° ± 45°; 277 electrodes, 21%) (Fig. S5). The functional significance of these phase relationships is explored in the following sections.

### Theta/High-Gamma phase amplitude coupling during speech

To understand how coherent theta oscillations might shape the timing of neuronal activity across the premotor speech network, we next asked whether the local theta phase modulates the magnitude of HG activity—an established marker of local population spiking activity (*53–55*)— in speech-responsive sites. Given the broad increase in theta phase coherence described above, even subtle theta-coupled enhancement of HG amplitude, when aggregated across coherently oscillating premotor sites, could give rise to a prominent pulsatile output signal. Much like crickets synchronizing their weak calls into a unified, pronounced pulsatile sound that can be heard from a distance, this coordination may serve to temporally align, integrate, and amplify neuronal signaling, focusing distributed premotor spiking within a theta-structured excitability window. In the context of articulatory control, such convergence is likely essential for broadcasting a unified, synergistic motor command to multiple articulators simultaneously.

To test this idea, we computed the phase-amplitude coupling (PAC) across speech-responsive sites using Tort’s Modulation Index (MI), as proposed by (*56*, *57*) (see Methods). We computed this index for each speech-responsive site individually and assessed statistical significance by comparing it to a surrogate distribution generated from 5,000 random circular shifts of the theta phase time series (Fig. S6; see Methods).

A widespread and statistically robust PAC was observed, with HG amplitude significantly modulated by local theta phase in 76.2% (975/1,280) of speech-responsive electrodes (P < 0.05, FDR-corrected; see Fig. S6 for details). Across electrodes, HG amplitude was highest near the theta trough (Fig. S6). It should be noted that this theta-rhythm modulation was often superimposed on slower, seconds-long HG activation profiles time-locked to sentence onset (see Fig. S1 for examples).

#### Quasi-rhythmic modulation of vocal tract kinematics during continuous speech

The large-scale theta phase coherence during speech, coupled with robust theta–HG PAC, suggests that a significant portion of speech-related premotor population activity is synchronized by the theta rhythm. At the mesoscopic scale, such in-phase oscillatory modulation can rapidly aggregate into a pulsatile neuronal signal at the output stations of the circuit, where motor commands are integrated and transmitted to the articulators. A direct prediction that follows is that the kinematics of vocal tract movements, encoded by the vSMC(*1*, *5*, *6*), should exhibit pulsatile dynamics coupled to the sensorimotor rhythm.

To test this prediction, we first analyzed electromagnetic midsagittal articulography (EMA) data from the Haskins Production Rate Comparison database (see Methods). These EMA recordings capture the kinematics of the upper vocal tract articulators (i.e., jaw, lips and tongue), with high temporal resolution (∼100 Hz), during the production of English sentences similar to those used in our study (Fig. 3A-B). Consistent with previous reports on rhythmic modulation of articulatory velocity during continuous speech (*58–60*), we found that the overall Articulatory Change (AC) during speech, namely, the sum of squared velocities (i.e., total speed) across the measured tract-variables, demonstrate rhythmic “pulses”, during which several articulators move together in synergy to form a constriction gesture (*5*, *58*) (Fig. 3B).

**Fig. 3.**
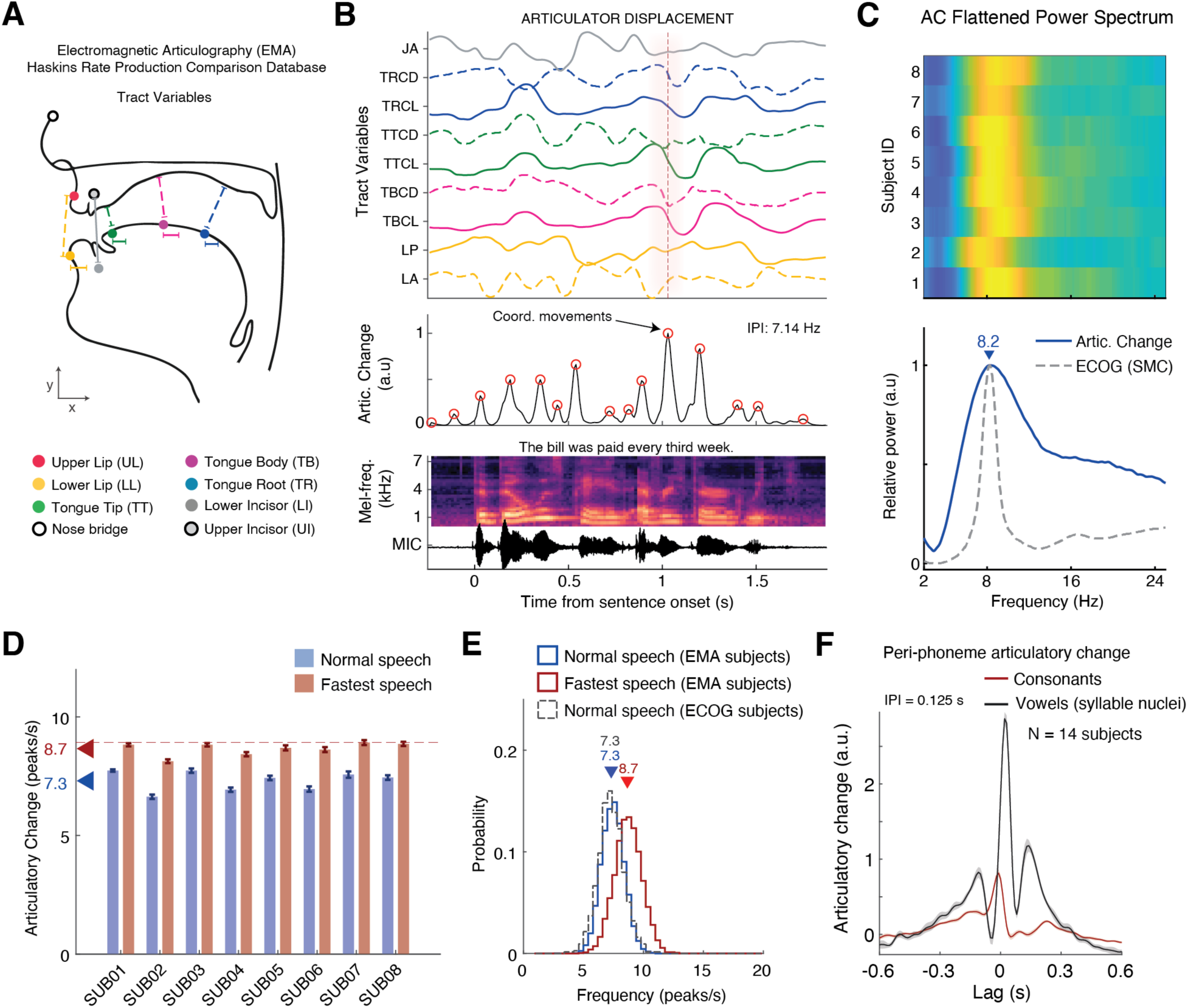
Quasi-rhythmic, pulse-like modulation of articulator velocity during continuous speech. (A) Example of articulatory kinematics for the jaw, lips and tongue monitored by Electromagnetic Midsagittal Articulography (EMA) during continuous speech production. (B) Top: Kinematic trajectories of the measured tract-variables including lip aperture and protrusion, tongue root, body and tip positions, and jaw angle (see Methods). Middle: We found semi-rhythmic temporal modulation of articulatory movements characterized by abrupt, pulse-like, changes in the sum of squared velocities across articulators. These brief pulses reflect coordinated movements associated with phonemes production (*58*, *59*). Bottom: Mel-spectrogram of produced acoustics. **(C)** Mean power spectrum showing a clear concentration of power around ∼8 Hz. The gray trace depicts the average power spectrum in SMC during articulation, derived from the ECoG subjects for comparison. **(D)** Analysis of the inter-peak intervals (IPI) of the AC time series during normal speech (∼4 syllables/s; blue bars) reveals a highly consistent median AC pulse rate across individuals (mean ± SE: 7.30 ± 0.15 peaks/s). When participants were instructed to speak as rapidly as possible without errors (red bars), the median AC pulse rate increased but remained centered below ∼9 Hz (8.66 ± 0.09 peaks/s). Error bars represent ± SEM across sentences. **(E)** Comparing the distribution of sentence-level AC pulse rates during normal speech in ECoG participants—where articulatory kinematics were estimated using AAI (dashed gray)—to those obtained from actual EMA measurements (blue) revealed no significant difference (P = 0.357, Wilcoxon rank-sum test; ECoG group: Med = 7.3, IQR = 1.4 pulses/s, N = 14; EMA group: Med = 7.3, IQR = 1.5 pulses/s, N = 8). For reference, the red distribution shows the upper bound of AC pulse rates, derived from EMA participants instructed to speak as quickly as possible. While this distribution shifts rightward, it remains centered within the theta frequency range (mean±SE: 8.66 ± 0.09 pulses/s). **(F)** Fitting a linear model to predict AC from produced phonemes (see Methods) revealed that the production of individual consonants (red) is associated with a single AC pulse. Phonemes forming the nucleus of a syllable (primarily vowels) elicit a larger AC pulse surrounded by secondary pulses, reflecting the preceding (onset) and following (coda) consonants that together with the nucleus form a syllable. Notably, the inter-peak interval of 0.125 s gives rise to a brief oscillatory pattern at 8 Hz.

In alignment with our prediction, the rate of these pulses during fluent speech consistently centered around 8 Hz. This was evident in both the flattened power spectrum of the AC time series (Fig. 3C) and the distribution of inter-peak intervals (Fig. 3D; see Methods). This quasi-rhythmic pulsatile behavior closely aligns with the mean peak frequency of the sensorimotor cortical rhythm across subjects (8.2 ± 0.7 Hz; Fig. 3C). Notably, when the EMA subjects were instructed to produce the same sentences as quickly as possible without errors, the rate of AC pulses increased but remained centered consistently below 9 Hz, i.e., within the sensorimotor theta range (median pulse rate averaged across subjects ± SE: normal speech [4 syl/s], 7.30 ± 0.15 pulses/s; fastest speech [5.8 syl/s], 8.66 ± 0.09 pulses/s; Fig. 3D).

While the EMA analysis revealed that speech movements exhibit rhythmic pulses in the theta range, these data alone cannot establish a direct relationship between cortical oscillations and vocal tract kinematics, as the neural and articulatory signals were recorded from different individuals. To determine whether AC pulses are temporally coupled to the ongoing sensorimotor theta rhythm, both signals—neural and kinematic—must be measured within the same subject. However, direct measurement of articulatory movements using EMA is impractical in ECoG subjects due to clinical and methodological constraints associated with bedside recordings. To overcome this limitation, we employed a deep learning-based acoustic-to-articulatory inversion (AAI) method (*5*, *61*), which infers the trajectories of the jaw, lips, and tongue directly from speech acoustics with high temporal precision. Fig. S7 shows example kinematic trajectories from a representative ECoG subject, along with the mean AC pulse rates across individual subjects.

To validate the fidelity of AAI-derived kinematics, we compared the AC pulse rates in ECoG subjects to those obtained from the EMA dataset during normal speech. The results showed no significant difference in median AC pulse rate between the two groups (ECoG vs. EMA: P = 0.36, Wilcoxon rank-sum test; Fig. 3E), confirming that the AAI approach preserves the fine-grained rhythmic structure of articulatory behavior during continuous speech.

### Articulatory Change in Relation to Phoneme and Syllable Production

To better understand the nature of the observed articulatory pulses, we next sought to characterize how the AC signal aligns with the elementary units of speech—phonemes and syllables. While prior work has characterized AC in relation to various articulatory landmarks (*58*), we extend this by isolating the fine-grained temporal profile of AC during the production of individual consonants versus vowels (i.e., syllable nuclei). Since the syllable is considered a fundamental unit of speech timing (*60*, *62–64*), and phoneme transitions are densely concentrated around its nucleus (*58*), we hypothesized that AC would exhibit a temporally structured sequence of pulses clustered around the vowel. As the most sonorous segment and the point of maximal vocal tract opening, the syllabic nucleus provides a natural temporal anchor for adjacent consonantal gestures (*65*), particularly during rapid consonant–vowel (CV) and vowel–consonant (VC) transitions. We therefore expected a prominent AC pulse aligned with the vowel, flanked by smaller pulses corresponding to the surrounding consonants.

To test this, we fit a linear multivariate temporal response function (mTRF) encoding model (*66*) to predict AC magnitude from phoneme onsets (see Methods). This approach allowed us to isolate the AC profile associated with individual consonantal gestures and compare it to the pattern emerging around the vowel.

Consistent with the idea that the syllabic nucleus acts as a temporal anchor, the model revealed a structured, quasi-rhythmic cluster of articulatory pulses centered around the vowel during syllable production. Fig. 3F plots the predicted AC magnitude in a −0.6 to +0.6 s window aligned to the onset of consonants (red) and vowels (black). Whereas individual consonants were tied to a single pulse, vowels were associated with a triplet of pulses corresponding to the syllable’s onset, nucleus, and coda. Notably, the mean inter-pulse interval in this triplet was 0.125 s—closely matching the 8.2 Hz sensorimotor theta rhythm. Supporting this, we also observed a strong correlation between AC pulse rates and syllable rates across subjects (R² = 0.94, P < 10⁻⁷; Fig. S7D), reinforcing the notion that AC pulses directly reflect the execution of articulatory gestures.

### Vocal-tract movements coupled to sensorimotor theta phase

If sensorimotor theta oscillations coordinate the timing of vocal tract movements during speech, then AC pulses should exhibit significant coupling to specific phases of the theta cycle. Since HG activity across the speech production network is strongly modulated by theta—peaking in unison around the trough (Fig. S6)—motor output is expected to reach local maxima shortly after these mesoscopic excitability windows. Accordingly, we predict that AC magnitude will transiently peak following these coordinated cortical activations.

To test this prediction, we binned the instantaneous theta phase into 24 bins (0 to 2π, 15° wide) and computed the average AC within each phase bin. If articulatory movements occur independently of theta phase, AC should be uniformly distributed across bins. To assess the coupling strength between theta phase and articulatory movements and determine statistical significance, we used Tort’s Modulation Index (*56*, *57*) and a non-parametric shuffling test, following the same approach used for the PAC analysis above (see Methods).

Fig. 4A-F presents the MI analysis for an example vSMC recording site, showing significant and consistent increases in AC coupled to the late phase of the theta cycle. Fig. 4D displays a single trial example from this site, demonstrating individual AC pulses that emerge consistently toward the end of concurrent theta cycles. Across 1,328 cortical speech-responsive electrodes in 14 subjects, we identified 204 sensorimotor sites (15%) with significant theta-AC coupling (P<0.05, FDR corrected; Fig. 4G). These electrodes formed a tight cluster along the central sulcus— spanning the pre- and post-central gyri and extending into posterior STG.

**Fig. 4.**
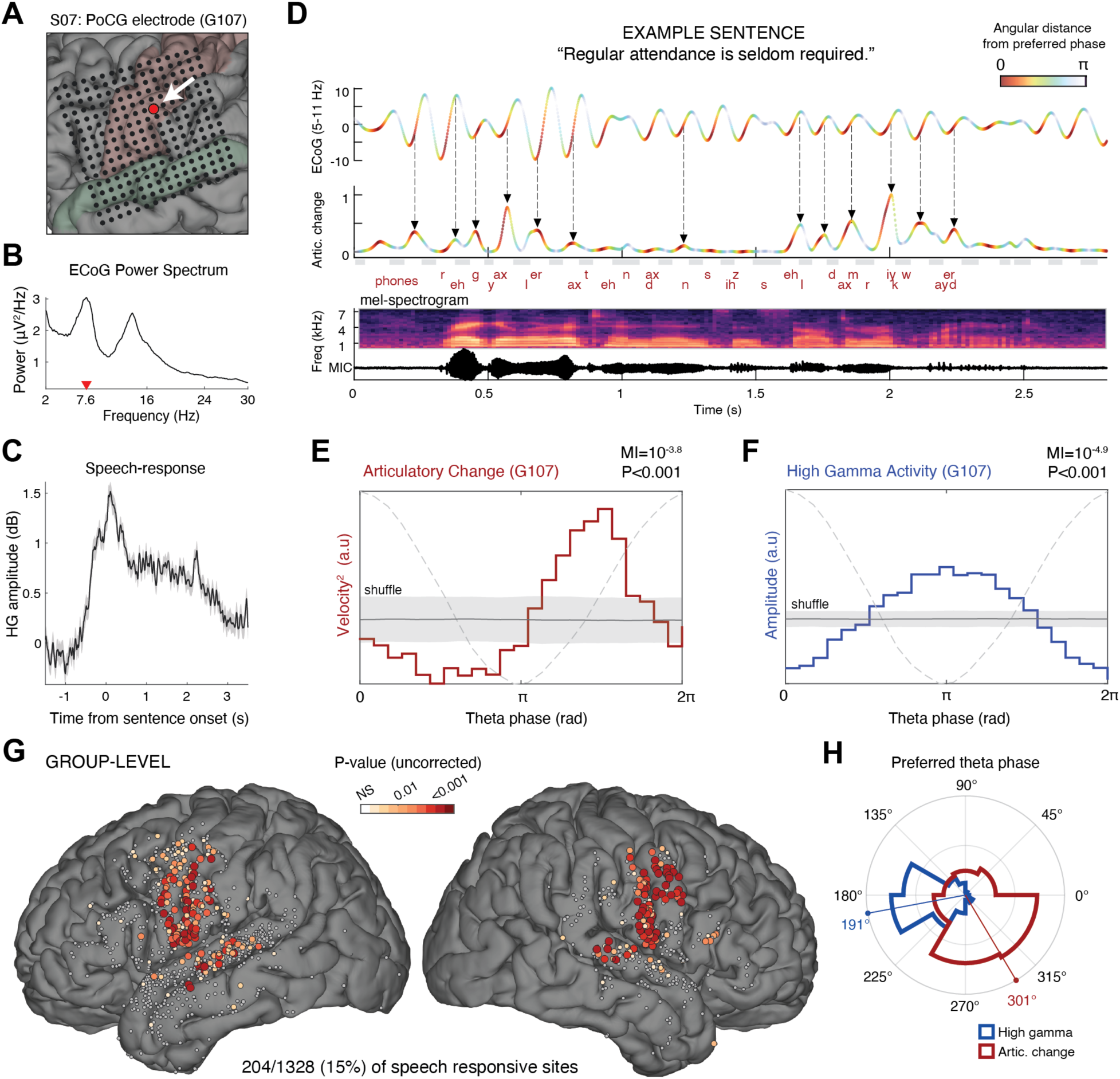
Articulatory change during continuous speech is coupled to the sensorimotor theta cycle. **(A)** Electrode’s anatomical location, **(B)** Power spectrum of the raw ECoG signal. Notice the spectral peak at 7.6 Hz (red triangle). **(C)** Mean HG response relative to sentence onset. **(D)** Example of a sensorimotor theta oscillation (top; 5-11 Hz bandpass filtered) and concurrent articulatory change (AC; middle) during the articulation of a representative sentence (produced phonemes, speech amplitude, and a mel-spectrogram are presented at the bottom). The theta rhythm was measured at a representative postcentral speech-responsive site described in panel a-c. The two timeseries are color-coded using the same colormap, indicating the angular difference from the preferred theta phase at this site, i.e., the phase at which articulatory change was maximal on average (see panel E). **(E-F)** We used Tort’s Modulation Index method (*56*) along with a shuffling procedure (see Methods) to statistically assess the coupling between theta phase and AC (E, red trace), and between theta phase and HG amplitude (F, blue trace). In this representative electrode, AC was significantly coupled to theta phase (MI=10^-3.8^, P<0.001), peaking a quarter cycle after the trough, while HG amplitude was maximal at the trough. Gray shaded area represent mean ±SD of the shuffled data; the dashed line represents the theta cycle. **(G)** Spatial distribution of recording sites where AC was significantly coupled to the theta rhythm (P<0.05, FDR corrected, 204/1328 (15%) of the speech responsive electrodes across 14 subjects). Note the anatomical specificity of the effect, clustering within SMC and the posterior STG. Non-significant electrodes are shown in light gray. **(H)** Polar histogram showing theta phase preferences for AC (red; P<10^-10^, Rayleigh test) and HG activity (blue; P<10^-44^, Rayleigh test). Only significant AC-theta coupled electrodes were included (N=204 sites). Notably, a consistent phase difference is observed between HG activity and A.C, with articulatory movements consistently preceded by HG activation within the theta cycle.

Notably, while the theta rhythm was observable and coherent across multiple sites within the speech production network, only this sensorimotor subset consistently translated the cyclic excitability windows into phase-locked motor output, highlighting its more direct role in articulatory execution.

In most sites (69%), the preferred phase—where AC was maximal—concentrated at the second half of the theta cycle (mean ± s.d.: 301°± 84°, P<10^-10^ Rayleigh’s test, N=204 electrodes; Fig. 4H and Fig. S8), approximately a quarter cycle after the HG amplitude enhancement found around the trough (preferred phase angular difference ± s.e., AC-versus-HG: 93° ± 5.8°, mean resultant length (MRL)=0.49, P<10^-15^ Rayleigh’s test; Fig. S8).

Interestingly, sites where AC was coupled to an earlier theta phase—specifically, before the trough—were consistently clustered in the dorsal portion of the ventral sensorimotor cortex (vSMC) (Fig. S8). This region has been implicated in articulatory motor planning (*67*), speech inhibition (*68*), and laryngeal control (*7*). These dorsal sites exhibited a theta rhythm that was nearly in anti-phase relative to more ventral “late-coupling” sites (mean phase ± SE relative to vSMC reference site: “early-coupling” electrodes (N = 26): –173° ± 24.5°; “late-coupling” electrodes (N = 90): 7° ± 4.6°; early vs. late: P < 0.001, Watson’s U² test; see Fig. S8 for details).

This bimodal distribution of theta phases is expected under the assumption that neighboring sites in the sensorimotor cortex are interconnected through both excitatory and inhibitory pathways (*69*). The existence of two anti-phase oscillators aligns closely with the coupled oscillator planning model (*65*)—a prominent model of speech timing positing that two theoretical oscillators, offset by 180°, govern the relative timing of articulatory gestures during syllable production. Within this framework, the in-phase/anti-phase theta pattern appears well suited to support both (i) transitions between the syllabic nucleus and coda elements (*65*), and (ii) the sequencing of complex consonantal gestures that require precisely timed transitions from full closure to release (*3*, *62*), as in oral plosives. Producing two such consonants in succession may leverage the 180° phase offset to schedule the second gesture half a cycle after the first, helping to prevent overlap—particularly when the same articulator is involved. Future research is needed to further elucidate the functional significance of this biphasic pattern across diverse articulatory contexts.

### Theta cycles orchestrate sequential premotor activations during articulation

The observation of systematic leading and lagging theta phases across sensorimotor sites suggests a sequential dorsoventral sweep of activation within each theta cycle. Specifically, early-phase sites would show theta-coupled HG activation first, followed by late-phase sites as the oscillatory cycle progresses. These theta-orchestrated activity sweeps naturally invite comparison to the phenomenon of “theta sequences” described in the rodent hippocampus, where sequential neural activity that unfolds over hundreds of milliseconds during behavior—e.g., as an animal moves through space—emerges in the same order but on a compressed timescale (∼125 ms) during individual theta cycles (*70*).

In these rodent studies, the greater the distance between two neurons’ place-field centers, the larger the difference in their preferred firing phases, indicating that spatial trajectories are mapped onto theta phase (*70*, *71*). Motivated by this analogy, we asked whether a similar relationship between behavioral and theta-compressed timescales exists for sensorimotor sequences encoding articulatory movements. Specifically, do electrodes that activate earlier relative to the onset of articulatory gestures also exhibit earlier relative theta phases?

To address this, we fitted a linear mTRF encoding model that predicts HG amplitude from articulatory kinematics (see methods). Crucially, this modeling approach enabled us to isolate the HG response associated with AC modulation while controlling for other kinematic features. It also allowed us to directly compare the peri-AC activation profile across electrodes with leading versus lagging theta phases. Importantly, since the encoding model operates exclusively in the time domain, it is entirely ’blind’ to the relative theta phase differences across sites. By comparing these time-domain peri-AC activation profiles with the independent phase relationships described earlier, we can assess whether phase order aligns with temporal order—that is, whether early-phase sites precede late-phase sites not only within the ∼125 ms window of a theta cycle, but also across the continuous, hundreds-of-milliseconds timescale of articulatory execution—resembling the theta sequences observed in rodents.

Across the electrodes that exhibited significant theta–movement coupling (P < 0.05, FDR corrected; N = 202 electrodes), we found a robust HG response time-locked to the onset of AC increase (Fig. 5A and Fig. S9). This peri-AC cortical activity was significantly stronger than the activation driven by any other kinematic feature, as shown by a timepoint-by-timepoint mixed-effects analysis (AC vs. all other features: P < 0.05, FDR corrected; Fig. 5A). These findings are consistent with our previous work demonstrating preferential encoding of both articulator speed (*6*) and multi-articulator gestures over single-articulator movements in the vSMC (*5*). Additionally, AC explained a significant proportion of unique variance across subjects (1 ± 0.09%, P < 0.0004, Wilcoxon signed-rank test; N = 14), confirming that it constitutes a key kinematic feature robustly encoded in the SMC.

**Fig. 5.**
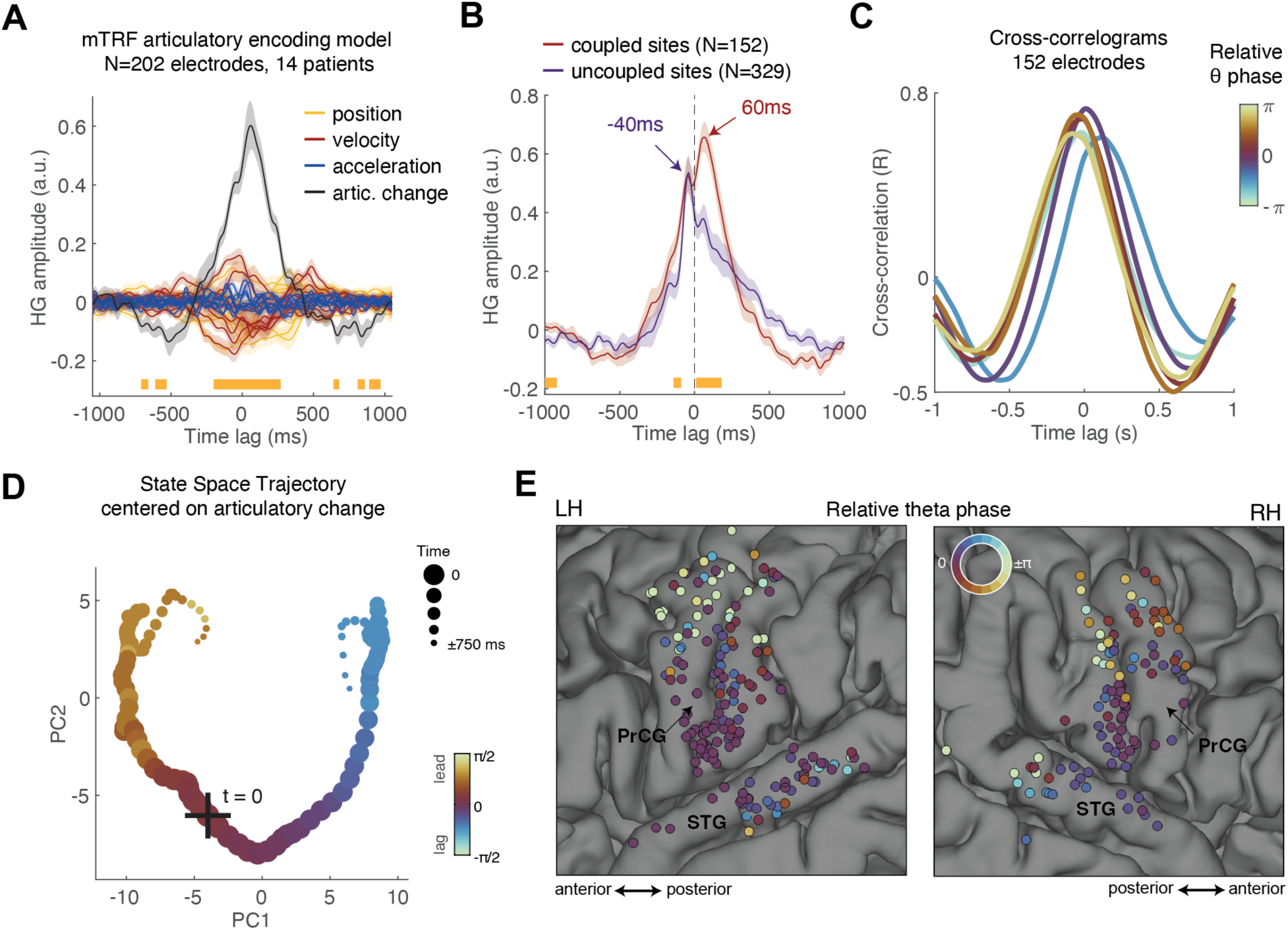
Sequential premotor cortical activations organized by theta phase. **(A)** HG activation profiles obtained from the mTRF articulatory encoding model, fitted to electrodes exhibiting significant theta-movement coupling (P < 0.05, FDR-corrected; N = 202 electrodes across 13 subjects). The weights corresponding to the Articulatory Change (AC) feature were significantly greater than those of any single-articulator feature (P < 0.05, FDR-corrected; timepoint-by-timepoint mixed-effects analysis comparing AC against all other features). Shaded error bars represent SE estimated from the mixed-effects model. **(B)** Peri-AC activation profiles averaged across ‘theta-coupled’ and ‘theta-uncoupled’ electrodes with significant unique variance explained by the AC feature. HG amplitude was significantly higher in ‘theta-coupled’ electrodes during the execution of articulatory movement (one-sided timepoint-by-timepoint mixed-effects analysis, P<0.05, FDR corrected). Shaded error bars represent SE estimated from the mixed-effects model. **(C)** Cross-correlograms between the peri-AC HG activation profile of each electrode and the average profile across all electrodes, computed separately for six relative-phase bins spanning –π to π. The cross-correlogram peaks shift systematically with the electrodes’ relative phase. **(D)** State-space trajectory computed by projecting the peri-AC activation patterns onto the first two principal components. The time points along the trajectory are color-coded based on the relative theta phase of the top 10% of electrodes with the strongest PC1 and PC2 weights at each time point. **(E)** Anatomical map of theta phase relative to the most ventral SMC site, including 283 electrodes in the SMC and STG exhibiting significant (P<0.05, uncorrected) theta-movement coupling across 14 subjects.

Next, we investigated whether electrodes exhibiting theta-movement coupling also demonstrated stronger AC encoding compared to those without such coupling. We pooled electrodes with significant unique variance explained by AC (P < 0.05, FDR corrected) into two groups: ‘theta-coupled’ electrodes (MI significance of P < 0.05; N = 157 electrodes across 14 subjects) and ‘theta-uncoupled’ electrodes (P > 0.1; N = 329 electrodes across 14 subjects). Comparing peri-AC activation profiles revealed that theta-coupled electrodes were more strongly driven by AC (P < 0.05, FDR corrected; timepoint-by-timepoint mixed-effects analysis of beta weights; Fig. 5B). In the ‘theta-coupled’ group, HG activity began increasing 300–400 ms before the AC and peaked 60 ms afterward, aligning with both premotor and execution-related activation. Conversely, in the ‘theta-uncoupled’ group, activity primarily anticipated the AC increase but culminated 40 ms before it, suggesting a predominant role in premotor preparation rather than the control of ongoing movement.

To test whether the relative theta-phase in ‘theta-coupled’ electrodes correlates with the onset of peri-AC response, we computed a cross-correlogram between each electrode’s peri-AC response and the grand-average response profile across all electrodes. We took the peak of this cross-correlogram as the electrode’s relative latency. We then assessed the correlation between these latencies and the electrodes’ theta phases, measured relative to each patient’s most ventral SMC site (see Methods).

This analysis revealed a robust circular–linear correlation between theta phase and the latency of peri-AC activation across the SMC (ρ = 0.41, P < 10⁻⁵; N = 152 electrodes), implicating theta phase in the temporal organization of vocal-tract control at the mesoscopic (multi-electrode, circuit-level-) scale. To visualize this relationship, we grouped electrodes into six phase bins spanning – π to π and averaged their cross-correlograms. As shown in Fig. 5C, the correlogram peak shifts progressively across phase-bins: electrodes with leading theta phase show earlier peri-AC activation. This relationship is further illustrated in Fig. 5D, which shows the state space trajectory of peri-AC activation in the SMC, with each point color-coded by the mean theta phase of the 10% most active electrodes (see Methods). Finally, Fig. 5E maps relative theta phase across ‘theta-coupled’ electrodes in SMC and STG, revealing a clear phase gradient along the SMC’s dorsoventral axis.

Lastly, to further quantify the relationship between the theta cycle and the continuous “behavioral” time, we computed the correlation between the latency differences across electrode pairs— estimated from cross-correlogram peaks—and their corresponding theta phase differences. Consistent with the preceding analyses, we found a highly significant correlation (*r* = 0.22, *P* < 10⁻¹⁶; Fig. S9), confirming that relative timing of peri-AC activation mirrors the phase structure of the ongoing theta rhythm.

### Consecutive theta cycles orchestrate successive movements during syllable production

Syllable production involves rapid transitions between successive constriction gestures associated with individual phonemes. Behaviorally, the temporal profile of AC relative to syllable nucleus suggests quasi-rhythmic transitions between gestures (Fig. 3F). Does the sensorimotor theta oscillation coordinate these transitions? To address this question directly, we epoched the ECOG and AC timeseries relative to syllable nucleus (i.e., vowels). In this analysis, we only included syllables that were well isolated in time, i.e., followed a period of at least 250 ms with no significant articulatory movement (see Methods). This resulted in a total of several hundred syllables per subject. For simplicity, the analysis was performed on a single sensorimotor site in each subject—the one that exhibited the strongest coupling between theta and AC (Fig. 6A-C).

**Fig. 6.**
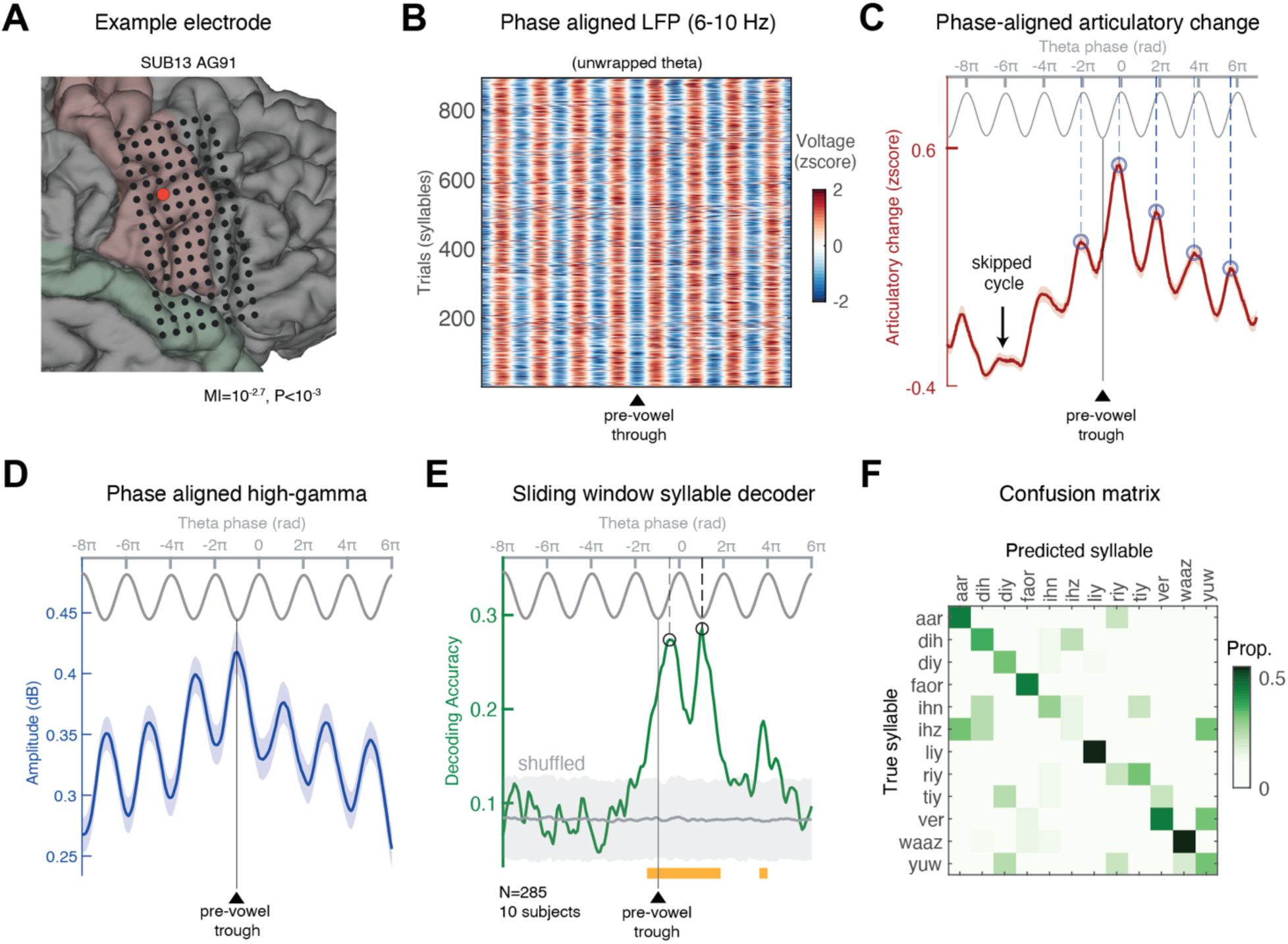
Consecutive theta cycles time articulatory movements during syllable production. **(A)** Anatomical location of a representative SMC site showing significant theta-movement coupling. **(B)** Each trial was aligned to the theta trough preceding the vowel (indicated by a black triangle). The theta phase was unwrapped and interpolated along a common phase axis spanning −9π to 7π (see Methods). Only syllables separated from the preceding one by at least 250 ms were included, allowing examination of whether theta–AC coupling persisted across brief pauses involving skipped cycles. **(C)** Articulatory change (AC) was interpolated along a uniform phase axis using the unwrapped instantaneous theta phase, as described in (B). The phase-aligned AC was then averaged across syllables. The analysis revealed consecutive AC pulses tightly locked to a specific phase of the theta cycle. In this electrode, AC peaked slightly before the theta peak (dashed blue vertical line). The arrow marks a cycle approximately 250 ms before syllable onset during which no articulatory movement occurred. Nevertheless, the subsequent movement remained coupled to the theta rhythm, consistently emerging at the preferred phase. **(D-F)** A sliding-window SVM syllable decoder was trained on the 12 most common syllables in the dataset, using 285 electrodes with significant theta-movement coupling, pooled from 10 subjects. (D) Phase-aligned HG amplitude, processed as described in (B), displayed pulsatile dynamics tightly coupled to theta troughs. (E) Decoding accuracy peaked during the two cycles overlapping with the produced syllable, slightly after the theta trough (peak phase: 4.07±0.15 rad). The gray-shaded area represents shuffled data (mean ± 95% CI), significant bins highlighted in orange. (F) Confusion matrix, computed by combining classifier outputs across the significant phase bins.

We then aligned each epoch to the theta trough closest to the syllable nucleus and extracted the unwrapped analytic theta phase for each epoch (see Methods). To ensure consistent phase alignment from beginning to end, we resampled the continuous phase series through linear interpolation onto a uniform phase axis ranging from [-9π to 7π]. The same interpolation was applied to the other signals, allowing us to average both the local population spiking activity (HG) and the concurrent AC across syllables while preserving their fine-grained alignment with the theta cycle (thus avoiding phase cancellation).

We found that the AC pulses associated with syllable production exhibited clear pulsatile dynamics that were tightly aligned with the ongoing theta phase, extending across several consecutive cycles. This is shown in Fig. 6C, which depicts the average AC in a representative site (see Fig. S10A-B for an additional example). Notably, because the analyzed syllables were preceded by brief articulatory pauses (see Methods), we were able to assess whether the first articulatory movement after such pauses “waited” for the next theta cycle. As shown in Fig. 6C, even after skipping an entire theta cycle, the subsequent movement consistently occurred at the preferred theta phase of that electrode.

For the group-level analysis, we centered the epochs on the preferred phase of each electrode (as determined by the MI analysis in Fig. 4) and averaged across subjects (Fig. S10C). For statistical testing, we generated surrogate data by circularly shifting the theta phases in each trial by a random amount 50 times (see Methods). Comparing the empirical and surrogate data revealed significant AC pulses emerging at the preferred phase across consecutive theta cycles—both in the cycle immediately preceding the syllable nucleus and in the cycles that precede or follow it. To confirm that this effect arises from the coupling of individual AC pulses, and not from a subset of particularly prominent movements, we also computed the group-level phase histogram (i.e., probability distribution) of AC peaks, which captures only the timing of the pulses rather than their amplitude. The results of this analysis, shown in Fig. S10E, indicate significant coupling of AC pulses to theta phase, irrespective of amplitude.

### Decoding syllabic content along the theta cycle

So far, we have shown strong coupling between theta phase, neuronal activity, and vocal tract movements. However, a key open question is whether the mesoscopic sensorimotor HG activity organized by the theta cycle actually represents speech content. Put differently, would a classifier trained to decode syllable identity from vSMC population activity exhibit performance that systematically varies with theta phase?

For the analysis we selected the 12 most common syllables in our dataset, i.e., those with a sufficient number of repetitions (N=10) to train a classifier (e.g., /ri/, /ɪz/, /fɔr/, /lɪ/, /ju/, etc.). We applied the phase-alignment procedure outlined above and centered all data epochs to the theta trough near the nucleus of the syllables. We then pooled together all electrodes that exhibited robust coupling (P<0.05, uncorrected) between theta and vocal-tract movement (n=285 across 10 subjects) and constructed a group-level matrix of HG activity during syllable production (electrodes ξ phase ξ trials). Crucially, like in the analysis above, this matrix aligned the activity in individual trials to a common phase axis ranging from [-9π to 7π], allowing us to link decoding performance to the instantaneous theta phase rather than time. Fig. 6D depicts the average HG amplitude across trials and electrodes, showing a strong pulsatile signal peaking at the trough of each theta cycle—as expected from the PAC described in Fig. S6.

Next, we trained a sliding-window Support Vector Machine (SVM) classifier (see Methods), with a 0.3-radian window, to decode syllable identity from HG activity patterns.

Fig. 6E–F presents the results of this analysis. Decoding performance was significantly above chance specifically during theta cycles that overlapped with syllable production (actual vs. shuffled labels: *P* < 0.05, FDR-corrected). Accuracy peaked shortly after the theta trough—i.e., when neuronal excitability is highest (Fig. 6D; phase of peak accuracy: 4.07±0.15 radians, ∼53° after the trough). When classifier outputs were combined across the significant phase bins (weighted by their mean accuracy), decoding accuracy reached 35.8% (chance = 8.33%). Fig. 6F displays the confusion matrix integrated across these significant bins. These findings underscore the role of theta oscillations in orchestrating mesoscopic, multi-site, sensorimotor activity that collectively represents the executed articulatory gesture during speech.

### Reduced Theta-Movement Coupling linked to Speech Errors

Our results identified strong coupling between theta and movement during fluent speech. If this coupling is important for orchestrating the complex motor sequences underlying speech, then we would expect it to be disrupted on trials where speech contained articulatory errors.

To test this, we analyzed utterances containing articulatory errors or speech repairs, focusing on electrodes showing significant theta–movement coupling. Subjects with fewer than 10 error trials were excluded. For each remaining participant, a control set of fluent trials was randomly sampled to match the error trials in syllable rate, duration, and sample size. This sampling was repeated 200 times to ensure stability, and the median of these resamples was used for analysis (Fig. S11).

Theta power did not significantly differ between fluent and error trials (*P* = 0.16, mixed-effects analysis; 117 sites, 7 subjects; Fig. S11A), suggesting that speech disruptions were not driven by changes in oscillatory power. Instead, it was the strength of theta–movement coupling (theta/AC MI) that was significantly reduced during speech errors (*P* = 0.001, mixed-effects analysis; Fig. S11B). Moreover, while the preferred theta phase was highly consistent across odd and even samples of fluent trials, error/repair trials exhibited greater variability in phase preference across electrodes. This was confirmed by significantly greater phase consistency across electrodes between two independent sets of fluent trials, compared to the consistency observed between fluent and error trials (resultant vector length: fluent = 0.99, errors/repairs = 0.73; P < 10⁻⁶, Fisher dispersion test; Fig. S11C-D). These findings suggest that speech errors are accompanied by a momentary disruption in the coupling between theta phase and articulatory movement, consistent with the idea that stable theta–movement coupling supports fluent speech.

### Theta Periodicity Predicts Individual Differences in Speech Rate

Lastly, the notion of sensorimotor theta as an internal timing mechanism for speech prompts a key question: are individual differences in speech rate driven by faster oscillations, or by more precise rhythmic timing? In other words, do fast speakers have a faster internal sensorimotor rhythm, or does their “metronome” tick more reliably, enabling more effective and precise coordination?

To address this, we analyzed the autocorrelation function (ACF) of the theta-band signal from the top five electrodes in each subject that showed robust theta–movement coupling. We first tested whether individual speech rate was associated with the frequency of the sensorimotor theta rhythm, but found no significant correlation (r = 0.18, *P* = 0.54, n.s.). However, when we assessed the periodicity of the oscillation—defined as the magnitude of the secondary ACF peak—we observed a striking relationship (Fig. S12A).

Subjects were divided into fast and slow speakers using a median split based on their mean syllable rate during the task. Fast speakers exhibited significantly stronger theta rhythmicity (*P* = 0.0006, rank-sum test; Fig. S12B), as indicated by more pronounced secondary ACF peaks (Fig. S12A–B), and greater theta-band power (P = 0.0012, rank-sum test). These results suggest that faster speech is not driven by a higher theta frequency, but by a more pronounced and temporally stable underlying rhythm.

Furthermore, theta rhythmicity correlated positively with speech rate during both sentence reading (Spearman’s ρ=0.79, P=0.001) and natural conversation (Spearman’s ρ=0.66, P=0.013; Fig. S12C–D). For the latter, we computed median articulation rates from spontaneous conversational utterances (mean length: 11 ± 3.1 words), excluding speech pauses. Together, these findings support the hypothesis that individual differences in speech rate are not driven by faster sensorimotor oscillations, but rather by greater stability and periodicity of the oscillations— consistent with the view of theta rhythm as an intrinsic timing mechanism that varies in strength and reliability across individuals.

## Discussion

In this study, we leveraged high-density electrocorticographic recordings in epilepsy patients to investigate the role of intrinsic neural oscillations in speech motor control. We identified a prominent theta rhythm (6–10 Hz) within the sensorimotor speech production network that supports the motor coordination required for fluent speech. Analysis of jaw, lip, and tongue movements during articulation revealed quasi-rhythmic, pulse-like modulations of articulator velocity, occurring ∼7-8 times per second on average. These motor pulses, reflecting rapid vocal-tract gestures, were tightly locked to specific phases of the cortical theta rhythm—with consecutive theta cycles timing successive phoneme-level gestures during syllable production. At the mesoscopic scale, the theta cycle orchestrated multi-site sensorimotor HG activation patterns, with syllabic identity optimally decodable shortly after theta troughs.

Our results thus demonstrate that mesoscopic representations of articulatory gestures are organized into precisely timed “pulses” of motor commands orchestrated by the theta cycle. Momentary disruptions in the coupling between theta-phase and articulatory movements predicted speech disfluencies, and individuals with stronger theta rhythmicity exhibited significantly faster speech rates.

These findings reveal a previously unrecognized role for theta oscillations in speech motor control, serving as a shared timing signal—an intrinsic temporal scaffold—that enables precisely timed, distributed control across multiple articulators moving in synergy. Such precise timing is essential for scheduling rapid transitions between articulatory gestures during fluent speech.

More broadly, our results highlight the critical role of intrinsic oscillations in achieving millisecond-scale coordination of distributed neuronal activity across mesoscopic brain circuits, providing direct empirical support for several existing theories on the function of neural oscillations (*12*, *16*, *36*, *39*, *43*, *72–78*).

Our findings also shed light on dynamics of phase coherence and PAC in cortical circuits (*79*). The co-occurrence of these phenomena suggests that, at the output stations of the oscillating circuit, spatially distributed activity may converge into a strong, distinct pulsatile signal. These mechanisms channel spatially distributed neuronal activity into discrete excitability windows aligned to specific oscillatory phases—a process particularly advantageous for coordinating muscle synergies with precise timing (*76*). Our demonstration of this mechanism in articulatory control parallels previous findings in the control of human hand and finger movements, where quasi-rhythmic kinematic pulses at ∼8 Hz have also been reported (*80*). Similarly, recent work in bats shows that wingbeat movements are highly rhythmic at ∼8 Hz (*81*). Together, these parallels suggest that the underlying premotor oscillatory mechanism may represent a general organizing principle for dexterous motor control, potentially extending beyond speech to other finely timed behaviors involving the hands, limbs, and orofacial effectors, and generalizing across species.

Prior work, dating back to early scalp and intracranial EEG studies (*82–84*), identified a rhythmic signal in the 8–13 Hz range, commonly referred to as the Rolandic mu rhythm or central alpha. This cortical rhythm has been traditionally studied in the context of motor tasks involving simple, discrete movements of the hands, feet, or tongue (*16*, *50*, *85–89*). In these early studies, sensorimotor oscillations often exhibited a time-locked power attenuation at movement onset, leading to their common characterization as the ’idle rhythm’ of the sensorimotor cortex—most prominent in the absence of voluntary movement (*16*, *90*). Later studies revealed more heterogeneous patterns, including both amplitude increases and decreases as well as inter-regional phase synchronization akin to what we observed—suggesting a more complex functional role with substantial task-dependent variability (*36*, *49*, *90*, *91*). Our findings extend this foundational work. By analyzing sensorimotor oscillations intracranially under the rapid and sustained motor demands of fluent speech production, we uncover a previously unrecognized role for these intrinsic sensorimotor rhythms in motor coordination. To distinguish this signal from the canonical mu rhythm—and to reflect its slightly lower frequency range (6–10 Hz) while maintaining alignment with related findings in animal models (*17*, *22*, *30*)—we referred to it as sensorimotor theta.

Our results reinforce the emerging view that theta rhythms are central to coordinating self-initiated movement and integrating it with cognitive processing—both in rodents (*17*, *20*, *22*, *24*, *30*, *92*, *93*) and non-human primates (*19*, *21*), underscoring the generalizability of these oscillatory dynamics across species and motor systems.

In line with this broader integration, we also find evidence of sensorimotor theta oscillations resonating within hippocampal circuits (Fig. 1E), suggesting a potential role in coordinating cortico-hippocampal communication during speech—a topic for future investigation. Relatedly, recent work (*94*) reported an increase in the coupling between neuronal spiking in the subthalamic nucleus and cortical theta rhythms during syllable production, supporting the idea that widespread theta phase coherence facilitates cortical–subcortical interactions during speech.

Critically, the intrinsic sensorimotor theta rhythm we report emerges independently of acoustic input. It is evident across diverse behavioral states, including silent rest, and maintains a stable central frequency despite large variations in speech syllable rate. A silent-miming control further dissociates this rhythm from acoustic feedback, underscoring a self-sustained premotor origin.

This finding provides a neurophysiological perspective that complements substantial EEG and MEG research implicating theta-band oscillations in auditory cortex during speech perception (*60*, *95*). In that work, theta activity within auditory cortex resets its phase to speech onsets and tracks the temporal envelope of the acoustic signal—processes linked to speech intelligibility (*96–98*). Perturbing the rhythmic structure of speech in the theta range likewise impairs intelligibility (*99*, *100*). Although the relative contributions of endogenous entrainment and evoked responses to oscillatory dynamics in auditory cortex remain an active area of investigation (*101*, *102*), converging evidence points to facilitatory effects on perception (*97*) and memory (*103*) when endogenous cortical rhythms align with the temporal structure of the acoustic speech signal.

Integrating auditory and motor findings, endogenous theta oscillations may differ in their sensitivity to acoustic input across regions of the speech cortex. Auditory-dominant areas can entrain to rhythmic structure in the incoming speech signal, whereas motor-dominant regions can oscillate independently of acoustics, supporting the intrinsic generation of distributed motor commands. Interactions between these cortical rhythms likely depend on cognitive state: during listening, envelope tracking in auditory cortex may facilitate alignment between perceptual and premotor processes, whereas during production, intrinsic premotor theta may align auditory regions with motor output to optimize acoustic feedback processing. Future work should clarify the relationship between the motor-related rhythm described here and the auditory rhythmicity found during perception, and determine how shared oscillatory dynamics across motor and auditory regions support feedback integration, speech comprehension, and memory (*103*, *104*).

The sensorimotor theta rhythm likely imposes fundamental constraints on the temporal limits of articulation. Recent work has identified a universal upper bound on speech production rate that closely aligns with the upper limit of the theta range (*60*). Coupé and colleagues (*105*), for instance, measured syllable rates across 17 languages and found values from 4.3 to 9.1 syllables/sec, averaging 6.63 syllables/sec, with virtually no language exceeding ∼7–8 syllables/sec.

Consistently, our EMA analysis showed that even during participants’ fastest speech, AC pulse rates remained below 9 pulses/sec (which depending on the precise proportion of open versus closed syllables (*58*) corresponds to ∼6-8 syllables/sec). These behavioral observations align well with our interpretation that the sensorimotor theta rhythm provides a rhythmic scaffold for coordinating speech movements, with the flexibility to skip cycles as needed (Fig. 6A–C) in order to generate output at rates below the theta range.

Finally, our findings also carry important clinical implications. The theta rhythm we identified may play a central role in speech disorders such as anarthria, apraxia of speech, stuttering, and various forms of aphasia—opening new avenues for therapeutic interventions targeting theta oscillations through neurofeedback or neuromodulation. Furthermore, our demonstration that speech decoding performance peaks around the theta trough has direct relevance for the development of speech neuroprostheses and brain-computer interfaces (*106*). This mesoscopic LFP rhythm opens a window into the “master clock” of distributed cortical processing—one that may be leveraged to optimize decoding accuracy and improve temporal precision in neural interfaces.

## Conclusion

Our study establishes sensorimotor theta oscillations as a core temporal organizer of articulatory gestures during fluent speech, opening new avenues for both theoretical inquiry and translational applications. This work sets a clear agenda for future investigations into rhythmic brain mechanisms underlying skilled, dexterous, and coordinated behaviors.

## Acknowledgments

We are grateful to the patients at UCSF for their kind participation and cooperation. We thank M. Leonard, I. Bhaya-Grossman, and L. Zhao for their valuable feedback on the manuscript. We thank S. Harper for guidance on articulatory phonology. We thank J. Hieronymus for helpful discussions on the results. We thank members of the Chang Lab for their assistance with data collection and their ongoing feedback throughout the project. We thank A. Fong for technical assistance with the recordings. We thank A. Silva and L. Zhao for guidance on AAI.

## Funding

National Institutes of Health grant U01NS117765 (EFC)

Jane Coffin Childs Memorial Fund for Medical Research (YN)

The Burroughs Wellcome Fund Career Award at the Scientific Interface (CASI) (YN) The Rothschild Fellowship (YN)

The Zuckerman STEM Leadership Program (YN).

## Author contributions

YN and EFC conceived the research and designed the experiment. YN, EFC and others collected the data. YN analyzed the data, LMF provided advice and feedback on the analysis, YN, LMF and EFC wrote and revised the manuscript. EFC supervised the project. The authors used AI-assisted language editing (ChatGPT, OpenAI) to improve language clarity. All scientific content, interpretations, and conclusions are the authors’ own.

## Data and materials availability

The data supporting the findings of this study will be made available upon reasonable request. Requests for materials should be addressed to EFC. The analysis code developed for this study will be made publicly available on GitHub following publication: https://github.com/itziknorman/Norman_et_al_2025_Sensorimotor_Theta

## Materials and Methods

### Participants

Seventeen participants (5 females, 12 males; age range: 22–60 years, M = 36.3) with medication-resistant epilepsy underwent chronic implantation of high-density subdural electrode arrays and depth electrodes as part of their evaluation for neurosurgical treatment (implanted hemisphere: 12 left, 5 right). Electrode locations were determined based on clinical requirements for each patient’s respective surgery. No participants had a history of cognitive deficits relevant to the aims of the present study. All participants were fluent in English. Participants provided written informed consent before participating in the studies. All procedures were approved by the University of California, San Francisco Institutional Review Board.

### Intracranial data acquisition

Electrocorticographic signals were acquired using subdural high-density grids (Integra or AdTech) with 1.17-mm-diameter exposed contacts and 4-mm inter-electrode spacing. The patients also had depth electrodes implanted in several subcortical regions. The voltage time-series (raw signal) from each electrode contact was amplified and digitized at a sampling rate of 3,051.7578 Hz using a pre-amplifier (PZ5, Tucker-Davis Technologies) and processed through a digital signal processor (RZ2, Tucker-Davis Technologies). During recordings, signals were referenced to the PZ5’s internal ground, which operated on battery power. This reference configuration proved remarkably stable, enabling low-noise, reliable recordings without the need to connect a head-mounted local reference electrode (e.g., a subgaleal electrode) to the preamplifier. Given the high signal quality, offline re-referencing was not required, allowing stable, artifact-free **monopolar** acquisition. Speech audio was recorded using a dynamic microphone (e845-S, Sennheiser), amplified by a microphone amplifier (MA3, Tucker-Davis Technologies), and digitized via the same RZ2 digital signal processor.

### Electrode anatomical localization

Pre-operative anatomical MRI and post-operative computed tomography (CT) scans were co-registered to determine electrode locations relative to the subject’s brain. The pial surface was reconstructed from the pre-operative T1-weighted and FLAIR MRI images using Freesurfer v7.3.2 (*107*). Electrode locations were manually labeled and snapped to the nearest point on the reconstructed cortical mesh. We used SUMA to resample and standardize the cortical surface of each subject (*108*, *109*), enabling visualization of electrodes from different subjects on a common SUMA-standardized cortical template (cvs_avg35_inMNI152). This approach preserves precise node-to-node correspondence across cortical meshes while maintaining the anatomical fidelity of each electrode’s position relative to the subject’s native gyri and sulci (*110*, *111*). Finally, electrodes were registered to the Desikan-Killiany gyral-based cortical atlas included in Freesurfer (*112*), allowing grouping by anatomical region. Cortical electrodes located more than 5 mm from the cortical ribbon were excluded from further analysis.

### Experimental paradigm: main task

Participants (N=14) read aloud 200 sentences from the MOCHA-TIMIT database (*113*) presented in four blocks of 50 sentences each, distributed across multiple days of the hospital stay. Three participants completed only two blocks (100 sentences). Within each block, sentences were shown one at a time on a laptop screen, and participants were instructed to read them aloud at their natural speaking rate following a visual go-cue. A random jitter of 1500 ± 500 ms was inserted between sentence presentation and the go-cue. The order of the sentences was randomized, and trials were advanced manually by the experimenter, allowing a few seconds of rest between consecutive sentences when needed. On average, utterance duration was 3.28 ± 0.61 s. Microphone recordings were obtained synchronously with the ECoG recordings. Trial onset and go-cue times were tagged using a photodiode and analog triggers sent on the audio channels. The MOCHA-TIMIT database, from which the sentences were drawn, is derived from the TIMIT corpus and designed to cover the full range of phonetic contexts in American English.

### Control tasks

Our study includes four key experimental controls: passive listening, silent rest, spontaneous speech production, and silent miming. All fourteen participants who performed the main reading task also completed a passive listening condition, during which they heard 225–600 sentences from the TIMIT acoustic-phonetic corpus (*114*), spoken by 286 male and 116 female speakers from various regions across the United States. Stimuli were presented through free-field speakers using custom MATLAB software on a Windows laptop. Sentences were delivered in pseudorandom order, with a 1 s silent interval between each.

Twelve participants additionally completed an autobiographical interview task, in which they were asked to recount personal experiences or memories; from these recordings, we extracted sentences spanning a broad range of natural, conversational utterances. Nine subjects also performed silent resting-state recordings lasting several minutes. These three conditions were compared to the main sentence-reading task (Fig. S3).

Finally, five subjects performed the silent-miming control task, which involved overtly producing a sentence, followed by a brief pause (∼1 s), and then silently miming the same sentence without producing sound (Fig. S2).

### Preprocessing and data analysis

Data analysis was performed in MATLAB 2024b (MathWorks Inc., Natick, MA) using EEGLAB v2021.1(*115*), the mTRF toolbox (*66*), Chronux v2.12 (*116*), CircStat toolbox (*117*) and custom-developed code. The raw iEEG signal was statistically inspected to identify and exclude noisy, disconnected, or non-functional channels. Specifically, channels whose voltage values, voltage derivatives, or RMS amplitudes in the top 1% (i.e., the 99th percentile) exceeded 5 standard deviations relative to other electrodes were flagged and subsequently reviewed via visual inspection in both time and frequency domains before exclusion. Preprocessing included notch filtering of the electrocorticographic signal to remove 60 Hz line noise and its harmonics at 120 Hz and 180 Hz, using a zero-lag, linear-phase Hamming windowed FIR band-stop filter (3 Hz wide), followed by resampling to 1000 Hz. Depending on the requirements of the specific analysis, the sampling rate was further downsampled to 500 Hz or 100 Hz.

### High-Gamma Activity

In this study, High-gamma (HG) activity—also known as High-Frequency Broadband (HFB) signal—was defined as the mean normalized power across frequencies from 60 to 160 Hz. Neural activity within this range is a well-established electrophysiological marker of local population firing (*118–120*). HG power was computed by: (1) band-pass filtering the ECoG signal in 20 Hz-wide bands (e.g., 60–80, 80–100, etc.) using zero-phase Hamming-windowed FIR filters (10 Hz transition width); (2) extracting the envelope of each narrow band signal by taking the absolute value of the analytic signal obtained from a Hilbert transform; (3) normalizing each amplitude time series by its mean; (4) averaging the normalized envelopes; and (5) multiplying the averaged time series by the mean amplitude across all bands, to restore voltage units. This normalization corrects for the 1/f spectral decay and yields a single broadband amplitude time series per electrode, reflecting local neuronal activity (*110*, *121*).

### Identification of speech responsive sites

We identified speech-responsive sites by comparing high-gamma (HG) amplitude during articulation versus the immediate pre-speech (–0.3 to –0.1 s) or post-speech (sentence end + 0.1 to 0.3 s) silent periods, using the Wilcoxon signed-rank test applied individually to each electrode. P-values from all electrodes were pooled to control the false discovery rate (FDR), and electrodes with FDR-corrected P < 0.05 in either comparison (relative to the pre- or post-speech window) were classified as speech-responsive. To quantify the magnitude of the response, we also computed the speech-response effect size using the standardized mean difference (Hedges’ g; see Fig. S1).

### Spectrograms

Depending on the analysis goal, time-frequency decomposition of the ECoG signal was performed using either the fast Fourier transform with a 1-second Hamming-tapered window or a Morlet wavelet transform, as implemented in EEGLAB (*115*). We used a window of 3 cycles at the lowest frequency (4 Hz) with a window scaling factor of 0.8 and a step size of 30–50 ms. Where relevant, spectrograms were normalized by dividing the power at each frequency by the geometric mean power computed over the pre-speech baseline window (−1 to 0 s), followed by conversion to decibels (10 × log₁₀) and baseline subtraction (*122*). For analyses focusing on the overall shape of the power spectrum (e.g., assessing the presence of a theta-band peak), rather than transient power fluctuations, power at each frequency was divided by the total power across all frequencies, yielding a spectrogram of relative power. To minimize the influence of transient spikes and electrical artifacts on spectral analysis, we inspected the raw signal from each electrode for voltage fluctuations exceeding 3σ relative to all other channels and excluded any trials in which such transients occurred.

### Phase Coherence Analyses

Event-related coherograms quantifying phase synchronization between two electrodes (a, b) across *n* trials were computed using the method described in EEGLAB’s ‘newcrossf.m’ function (*123*):

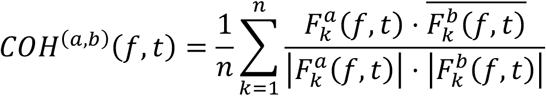

where 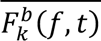 denotes the complex conjugate of 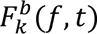. The normalization factor in the denominator ensures that only the relative phase relationship between the two spectral estimates is considered across trials, independent of differences in power. Coherence magnitude ranges from 0 to 1, with 0 indicating a complete absence of phase-synchronization at frequency f in the time window centered at time t, and 1 indicates perfect phase locking. The phase difference between electrodes can then be computed by taking the angle of the complex-valued coherence estimate (across 𝑛 trials). For consistency across subjects, depth electrodes were excluded from the analysis. To minimize potential contributions from passive volume conduction, pairwise coherence was computed only between speech-responsive grid electrodes separated by at least 15 mm (see Fig. S5A).

### Relative theta phase across speech-responsive electrodes

To assess systematic theta-phase differences among speech-responsive electrodes during continuous speech (Fig. S5, S8; Fig. 5) we filtered the raw ECoG signal in the theta range (5– 11 Hz) using a zero-lag, linear-phase FIR bandpass filter with a 2 Hz roll-off (Hamming window) and extracted the instantaneous phase using the Hilbert transform. A reference site in the most ventral vSMC was selected based on anatomical location and theta power. For each speech-responsive electrode, relative theta phase was quantified as the mean phase difference from this reference site across artifact-free speech intervals (see above).

### Power Spectral Density Analyses

To obtain a robust estimate of the power spectral density (PSD) of the raw ECoG signals, we employed the multitaper method (*124*, *125*), as implemented in Chronux 2.12 (*116*), an open-source MATLAB toolbox. The multitaper approach reduces the variance of spectral estimates by applying multiple orthogonal windowing functions (Slepian tapers) to the data. This produces a set of independent spectral estimates, which are then averaged to yield a more stable and reliable PSD estimate. The continuous intracranial ECoG signal recorded during the task was segmented into 6-second epochs aligned to speech onset (−1 to +5 s). Spectral estimates were computed using five Slepian tapers, yielding a frequency resolution of 1 Hz. Prior to analysis, all data segments were demeaned and zero-padded to a length of 4,096 time points (∼8 s).

### Detection of Oscillatory Peaks in the Power Spectrum Peak

To detect spectral peaks and assess their prominence, we applied two complementary methods. In Fig. 1C, we computed the power spectral density after first standardizing each voltage time series by rescaling its values between the 1st and 99th percentiles. This normalization ensured a comparable dynamic range across electrodes, allowing consistent comparison and visualization of spectral profiles while minimizing bias from inter-electrode amplitude differences. The multitaper power spectrum was then computed on the rescaled signals. To isolate spectral peaks, we removed the 1/f background by fitting a linear regression to the 2–30 Hz range and removing the fitted trend. The resulting spectrum was z-scored, smoothed using a Savitzky–Golay filter (2nd-order polynomial, 1 Hz window; ‘sgolayfilt.m’), and subjected to peak detection. Peak prominence was defined as the height of a peak relative to its surrounding baseline, expressed in z-scored power values.

In all other analyses of spectral peaks, we applied a parametric power spectrum decomposition using the FOOOF algorithm (*48*) which transforms power to a log-scale, separates the aperiodic 1/f component from the oscillatory components and then applies peak detection on the 1/f corrected spectra. We used the default FOOOF parameters (2–30 Hz frequency range; peak threshold = 2.5) and excluded peaks with widths > 6 Hz. When comparing across different tasks (e.g., Fig. S3), we applied the voltage-rescaling procedure described above prior to power spectrum estimation and FOOOF analysis. When comparing across speech rates (Fig. 2A-C), we skipped this step, as the comparison is done within electrode.

### Quantifying Phase-Amplitude Coupling (PAC)

Phase-amplitude coupling (PAC) was measured using Tort’s Modulation Index (MI) approach (*56*, *57*). Instantaneous theta phase was extracted via the Hilbert transform and divided into 24 equally spaced bins spanning the full [0, 2π] range. High-gamma (HG) amplitude was averaged within each phase bin during speech intervals, excluding pre- and post-utterance silent periods. The resulting phase–amplitude distribution was compared to a uniform distribution using Kullback–Leibler divergence to quantify the degree of modulation. This yielded an MI value reflecting the extent to which HG amplitude was modulated by theta phase. PAC was computed separately for each electrode, and statistical significance was assessed by comparing the observed MI to a surrogate distribution generated from 5,000 random circular shifts of the theta-phase time series. To quantify coupling between theta phase and articulatory change (AC), we applied the same procedure, this time using the AC time series. To ensure comparability across electrodes and subjects, both HG and AC time series were rescaled to the [0, 1] range prior to computing the modulation index, allowing the analysis to focus on phase coupling independent of absolute amplitude differences. To generate the comodulogram shown in Fig. S6, we computed the MI across multiple narrowband frequency pairs: phase frequencies were filtered using a fixed 4 Hz bandwidth in 1 Hz steps from 4 to 20 Hz, and amplitude frequencies were log-spaced from 20 to 160 Hz, with bandwidths increasing from 4 Hz at the lowest frequency to 40 Hz at the highest. For consistency across subjects, only speech-responsive electrodes located on the cortical grid were included in this analysis; depth electrodes were excluded.

### Acoustic-to-Articulatory Inversion (AAI) and Tract Variables

Many critical vocal tract movements involved in speech production are not externally visible and cannot be easily monitored. Capturing the kinematic trajectories of articulators such as the jaw, lips, and tongue requires imaging techniques capable of dynamically tracking both external and internal articulatory motion during continuous speech. One such method is electromagnetic midsagittal articulography (EMA), which uses sensors placed on key points of the vocal tract to monitor their movement in real time within a magnetic field during fluent speech.

However, because simultaneous acquisition of EMA and ECoG data is not practically feasible, we employed a deep-learning-based method to infer articulatory dynamics directly from the acoustic signals recorded via microphone (*5*, *61*). This state-of-the-art acoustic-to-articulatory inversion (AAI) technique operates in the *tract-variable* space—a set of parameters derived from the trajectories of the tracked EMA sensors, that quantify the degree and location of vocal-tract constrictions. These variables capture key geometric distances along the vocal tract that define each constriction gesture. This representation yields a speaker-independent, gesture-relevant model of articulatory behavior (*3*, *126*) and enables accurate AAI-based reconstruction of articulatory kinematics that generalizes well to unseen speakers (*61*).

Using the AAI technique, we monitored the kinematic trajectories of 9 tract-variables at 100 Hz temporal resolution: LA, lip aperture; LP, lip protrusion; TBCL, tongue body constriction location; TBCD, tongue body constriction degree; TTCL, tongue tip constriction location; TTCD, tongue tip constriction degree; TRCL, tongue root constriction location; TRCD, tongue root constriction degree; and JA, jaw angle.

The geometric transformations used to compute tract-variables from EMA sensor positions are detailed in (*61*, *127*). We applied these transformations to raw EMA measurements from the Haskins Rate Production Comparison (HPRC) database (*128*) and derived tract-variable trajectories representing ground-truth articulatory kinematics. These were analyzed in parallel with the AAI-inferred trajectories obtained from our intracranially monitored ECoG participants (Fig. 3). This comparison served to validate the AAI reconstructions and to ensure that our characterization of vocal tract kinematics using the Articulatory Change (AC) feature was not biased by potential inaccuracies in the inversion process.

### Articulatory Change

To capture the dynamics of articulatory change over time—that is, the temporal modulation function of the articulatory process—we adapted a method recently proposed by Goldstein (*58*) to our nine EMA-derived tract-variables. This method quantifies articulatory change as the sum of squared velocities across tract-variables:

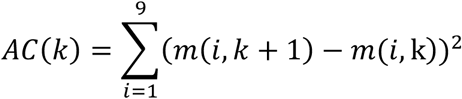

where 𝑚(𝑖, 𝑘) denotes the value of the 𝑖-th tract variable at frame 𝑘. This univariate signal reflects the instantaneous magnitude of articulatory movement across the vocal tract and serves as a proxy for the temporal modulation of articulatory gestures during continuous speech. To align the AC time series with the sampling rate of the ECoG signal, we first computed the tract-variable derivatives at its original 100 Hz sampling rate, then upsampled the resulting time series to the target resolution using spline interpolation.

### Articulatory Change Pulse Rate

To compute the rate of AC pulses during continuous speech while maintaining consistent detection criteria across subjects, we first applied peak detection to sentence-level AC traces from the EMA dataset. Utterances with syllable rates below the 1st percentile or above the 99th percentile across all sentences, or those containing articulatory pauses longer than 0.5 seconds, were considered outliers and excluded. AC pulses were then identified using MATLAB’s findpeaks.m function, with a minimum peak distance of 20 ms. From the detected peaks, we derived group-level distributions of peak prominence and width values, pooled across both normal and fast speech. The 1st percentile of each distribution was selected as the threshold for subsequent peak detection. These empirically derived thresholds were then applied to both the EMA and ECoG datasets, ensuring uniform and data-driven pulse detection across all subjects and conditions. In all subsequent analyses, excessively slow sentences—defined as those with syllable rates falling below the first quartile minus 1.5 times the interquartile range, or those containing articulatory pauses longer than 0.5 seconds—were excluded.

### Phonetic and phonological transcription

Transcriptions of the recorded speech acoustics began with automatic speech-to-text transcription using Adobe Premiere Pro 5, followed by manual correction at the word level, ensuring that the transcript reflected the vocalization that the participant actually produced. Using these sentence-level transcriptions and the corresponding acoustic utterances, we performed sub-phonetic alignment following previously described methods (*129*, *130*), and extracted the onsets of individual phonemes, including vowels constituting syllabic nuclei. To ensure a stable and standardized method for counting syllables per utterance, we used the Datamuse API (https://www.datamuse.com/api) to retrieve dictionary-based syllabification for each produced word or its closest phonological match, based on American English pronunciation.

### Speech errors

During transcription, utterances containing speech errors that resulted in significant fluency disruptions or were immediately followed by a speech repair were logged and excluded from subsequent analyses. These error trials were then specifically analyzed in Fig. S11. Typical examples included substitutions or mid-word stopping followed by immediate repair, such as: *“Curiosity and media(*) mediocrity seldom coexist”* or *“Publice(*) publicity and notoriety go hand in hand”.* The error trial rate across participants was 7.5 ± 5.8% of sentences.

### Multivariate temporal response function (mTRF) encoding models

We used the mTRF toolbox (*66*) to fit multivariate encoding models using ridge regression (*5*, *131*, *132*). For a detailed mathematical description of the fitting procedure, see the original reference of the toolbox. Regularization strength (λ) was optimized using 10-fold cross-validation on a training set comprising 90% of the data. Model performance was evaluated on the remaining 10% (held-out test set) by computing the Person’s correlation (r) between the predicted and actual signals.

This framework was applied in two analyses. First, to recover the temporal profile of articulatory change (AC) during the production of individual phonemes and syllables (Fig. 3F), we modeled the AC time series as a linear sum of overlapping event-related responses time-locked to phoneme onsets. Phonemes were grouped into six classes: stops, fricatives, affricates, nasals, liquids, and syllabics. Both the phonemic predictors and AC responses were z-scored within each trial. The model was trained on data resampled to 100 Hz, with time lags from –1250 to +1250 ms. On average, the fitted models—validated on a held-out test set—achieved an r value of 0.37 ± 0.05 (mean ± s.d) up to r=0.5.

In the second analysis, we modeled the HG amplitude time series at each electrode as a weighted sum of articulatory kinematic features over time. These features—derived from AAI-extracted tract variables (Fig. 3)—captured movements of the jaw, lips, and tongue. The model included the instantaneous position of the articulators (constriction degree and location; 9 features), their velocity (first derivative; 9 features), acceleration (second derivative; 9 features), and their overall speed—i.e., the Articulatory Change (AC) time series—totaling 28 features. To ensure comparability across features and trials, both the kinematic predictors and HG responses were z-scored within each trial (i.e., centered and normalized). The model was trained on data resampled to 100 Hz, with time lags from –1050 to +1050 ms. In electrodes where theta phase was significantly coupled to AC, the fitted models—validated on a held-out test set—showed r value of 0.22 ± 0.13 (mean ± s.d), up to r=0.54, consistent with our previously reported estimates in vSMC (*5*). To determine whether the AC feature itself accounted for a significant amount of unique variance, we repeated the modeling procedure using a temporally shuffled version of the AC predictor. Model performance (r²) was then compared between the original and shuffled models across 200 iterations (see Fig. S9A). Statistical significance was assessed by calculating the proportion of iterations in which the shuffled model’s performance equaled or exceeded that of the original model.

### State space analysis

To investigate the relationship between theta phase and the mesoscopic SMC activation elicited during articulatory movements, we projected multi-electrode HG activity patterns onto a two-dimensional state space (*1*). This projection was computed by applying principal component analysis (PCA) to z-scored peri-AC HG activation profiles derived from the mTRF encoding model, including only SMC electrodes that showed significant theta-movement coupling. The first ten components accounted for 64.6% of the total variance. HG activity was then mapped onto the first two principal components (explaining 33% of the variance) and color-coded according to the circular mean of the relative theta phases of the most active electrodes at each point along the trajectory—defined as those with the top 10% of PC_1_ and PC_2_ scores.

### Phase-aligned averaging of peri-syllable articulatory change

To investigate the relationship between consecutive theta cycles and AC during sequential gestures grouped within a syllable, we developed a method that enables averaging across events of interest—i.e., syllabic nuclei—while preserving multi-cycle phase alignment between the ongoing theta oscillation and the concurrent AC signal (Fig. 6A, Fig. S10). Theta-band–filtered ECoG and AC time series were epoched from –750 to 750 ms relative to the syllable nucleus (i.e., vowel onset). Only syllables that were temporally isolated—preceded by at least 250 ms of articulatory pause—were included. Articulatory pauses were defined as periods without detectable AC pulses, identified using empirically determined peak prominence and width thresholds, based on actual EMA recordings from the Haskins Production Rate Comparison (HPRC) database (see the *Articulatory Change Pulse Rate* subsection for details). Syllables with extreme durations (outside the 1st and 99th percentiles) were excluded, yielding several hundred valid syllables per subject. For simplicity, analyses were restricted to a single sensorimotor electrode per subject— specifically, the one exhibiting the strongest theta–AC coupling.

To preserve phase alignment across syllables, each epoch was aligned to the theta trough closest to the syllable nucleus, and the unwrapped analytic theta phase was extracted. Continuous phase series were then resampled via linear interpolation onto a uniform phase axis spanning eight consecutive theta cycles (from –9π to 7π relative to the nucleus-aligned trough). The same interpolation was applied to the AC signal, enabling across-syllable averaging while maintaining precise phase alignment with the theta cycle—thus avoiding phase mixing or cancellation.

In the group-level analysis, phase-aligned AC profiles were averaged across subjects after re-aligning each electrode to the phase of maximal AC in that electrode, as determined in Fig. 4E. To assess the statistical significance of AC modulation by theta phase, the actual observed pattern was compared to a surrogate distribution generated by circularly shifting the theta phase by a random amount and repeating the full analysis 50 times. Because both real and shuffled analyses centered epochs on the trough closest to the syllable nucleus, this procedure preserved slower dynamics associated with syllable production while disrupting only the fine-grained theta phase relationship. Contrasting the observed and shuffled profiles thus isolated the specific contribution of theta phase to articulatory change during syllables. We quantified this contrast as the percent change between the observed and shuffled profiles for each electrode and tested it against zero (N=14, one electrode per subject) using Wilcoxon signed-rank tests (Fig S10). P-values were corrected for multiple comparisons across phase bins using FDR.

### Phase-aligned syllable decoding analysis

To determine whether articulatory information encoded at the mesoscopic scale emerges transiently at specific phases of the theta cycle, we trained a sliding-window syllable decoder that advanced along the unwrapped theta phase axis, rather than time. We selected the 12 most common syllables in our dataset (e.g., /ri/, /ɪz/, /fɔr/, /lɪ/, /ju/)—defined as those with at least 10 repetitions across sentences, sufficient to train a classifier. The HG time series was aligned across trials using the procedure described above, centering each data epoch on the theta trough closest to the syllable nucleus. Electrodes exhibiting robust theta–movement coupling (285 sites across 10 subjects) were pooled to construct a group-level matrix of HG activity during syllable production (electrodes × phase × trials). As in the analysis in Fig 6A-C, activity in each trial was aligned to a common phase axis spanning eight consecutive theta cycles (from –9π to 7π relative to the nucleus-aligned trough). This phase alignment enabled decoding performance to be tracked as a function of instantaneous theta phase, rather than time, across multiple consecutive cycles (Fig. 6D-F).

For the decoding analysis, phase-aligned HG activity patterns were z-scored across trials and re-binned into 0.8-radian-wide phase bins with a 0.3-radian step size, ensuring a fixed phase resolution across electrodes. To account for the systematic phase differences observed along the dorsoventral axis of the SMC, electrodes were re-aligned to the phase of articulatory movement (i.e., the phase of maximal articulatory change (AC) in each electrode, as determined by the analysis described in Fig. 4E). This step ensured that signals were not only phase-aligned across trials within each electrode—but also centered on a common physiological reference across electrodes: the phase of maximal articulatory movement nearest the syllable nucleus.

A multi-class SVM classifier was trained to decode syllable identity from these phase-aligned HG activity patterns using 5-fold stratified cross-validation, repeated independently at each phase bin (using MATLAB functions fitcecoc.m and cvpartition.m). Decoding accuracy was evaluated on held-out test folds, and the analysis was repeated 25 times with different initializations to obtain a stable estimate of decoding accuracy. To assess statistical significance, a null distribution was generated by repeating the entire decoding procedure 500 times using shuffled trial labels. Crucially, the same permutation was applied across all phase bins within each shuffle iteration, preserving temporal structure while disrupting the syllable label mapping. This allowed fair comparison between real and shuffled decoding performance. Accuracy was smoothed using a 5-point Savitzky–Golay filter (2nd-order polynomial). P-values were computed as the proportion of shuffles in which accuracy exceeded or equaled the mean accuracy of the real data and were corrected for multiple comparisons using FDR.

### Statistical tests

All statistical analyses were performed in MATLAB. Unless otherwise noted, pairwise comparisons were conducted using two-sided Wilcoxon signed-rank or rank-sum tests. For circular data, we used the CircStat toolbox (*117*). The unit of analysis was typically individual subjects, or electrodes nested within subject when using mixed-effects analyses. In some cases, as stated in the main text, the unit was individual electrodes or activity patterns across pooled electrodes (as in the syllable decoding analysis). Resampling tests were performed using custom MATLAB scripts based on previously published routines (*133*, *134*), or with algorithms from the Mass Univariate ERP Toolbox (*135*). Multiple comparisons across electrodes or time bins were corrected using false discovery rate (FDR) adjustment (*136*). Data collection was conducted blind to experimental conditions; data analysis was not. No statistical methods were used to predetermine sample sizes, but sample sizes were consistent with those commonly used in the field (*68*, *111*).

### Mixed effects analysis

Mixed-effects analyses were conducted in MATLAB using the ‘fitlme.m’ function. Models included the relevant fixed effects and a random intercept for ‘Subject’, with ‘Electrode’ nested within ‘Subject’ (or ‘Electrode-pair’ nested within ‘Subject’, as in the pairwise coherence analysis). For example, to test for differences in central frequency across speech rates (fixed factor *condition* with three levels—slow, medium, fast), we used the following mixed-effects model:

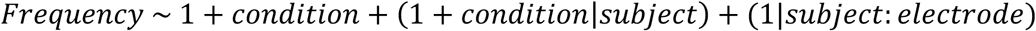

This nesting of electrodes within subjects accounts for the hierarchical structure of the data— specifically, that each participant contributed multiple electrodes (or electrode pairs) to the analysis, and that observations from the same subject or electrode are not statistically independent. It also captures variability arising from the fact that participants contributed different numbers of electrodes. Random slopes were included when justified by the data structure, provided they did not lead to over-parameterization or model non-identifiability. Main effects were tested using Type III ANOVA implemented in ‘fitlme.m’. Degrees of freedom were estimated using the Satterthwaite approximation. For analyses involving timepoint-by-timepoint mixed-effects models, *p*-values were computed individually at each time point and corrected for multiple comparisons using FDR.

---

**Fig. S1.**
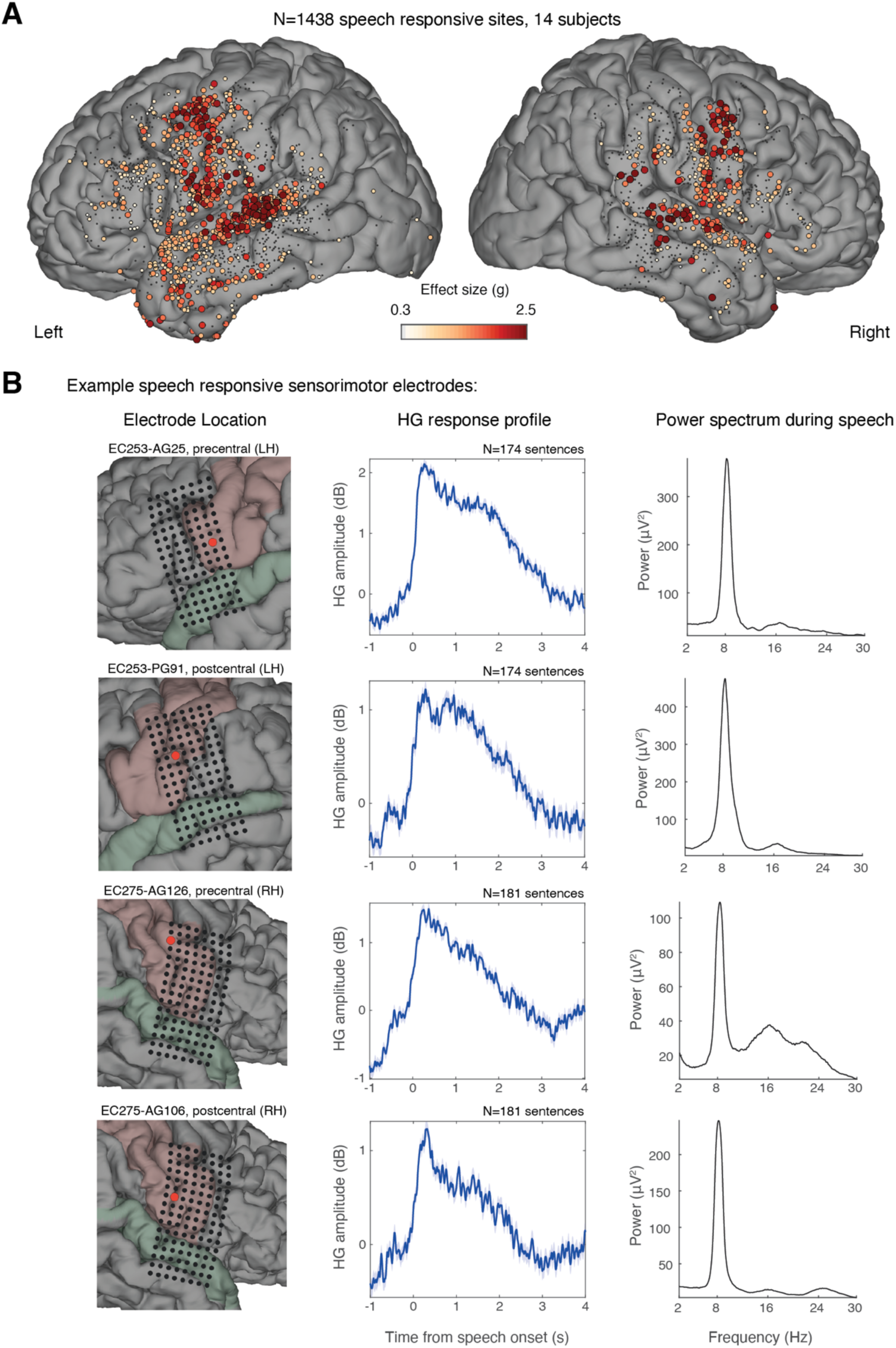
Overview of speech responsive electrodes. **(A)** Anatomical distribution of speech responsive electrodes. Out of 3,599 recording sites analyzed, 1,438 were classified as speech-responsive, showing a marked increase in HG amplitude during speech compared to the inter-trail silent periods (P < 0.05, FDR corrected). The spatial distribution of these speech-responsive sites highlights the key cortical regions involved in speech production, including the sensorimotor cortex (SMC), superior temporal gyrus (STG), supramarginal gyrus (SMG), and additional sites in the prefrontal and temporal cortices. We used this gross functional classification of electrodes to focus subsequent analyses on speech-responsive sites, effectively excluding task-irrelevant sites. **(B)** Example of speech-responsive SMC electrodes from two patients. The left panels show the anatomical location of the electrode, the middle panels display HG activity (60-160 Hz) during speech, and the right panels present the multi-taper power spectrum computed over an interval of -1 to 5 s relative to sentence onset, showing a prominent 8 Hz LFP oscillation occurring alongside the HG activity associated with speech production.

**Figure S2.**
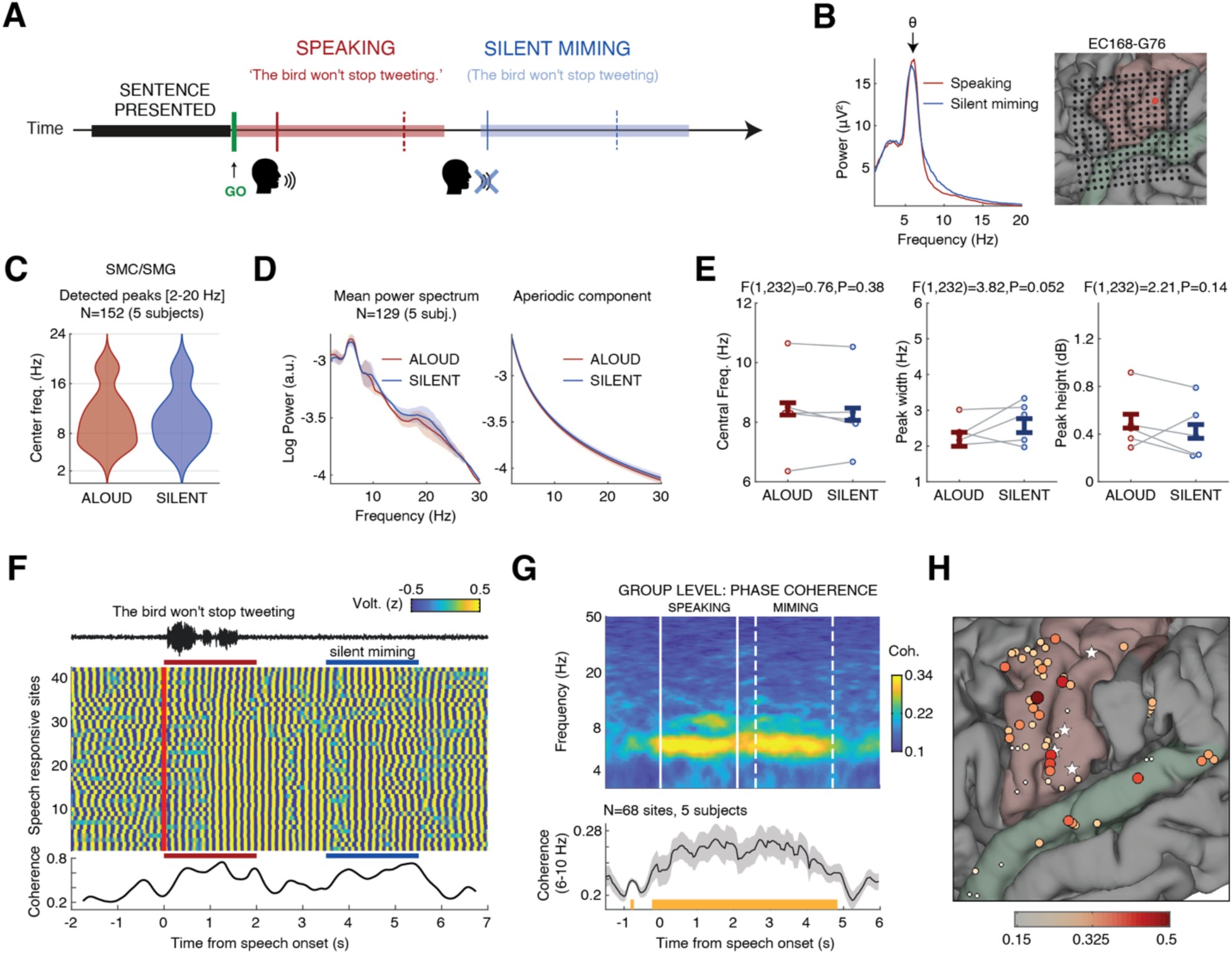
Coherent sensorimotor theta rhythm persists during both overt speech and silent miming. (A) Five subjects performed a control task that involved overtly producing a sentence, followed by a brief pause (∼1 s), and then silently miming the same sentence. (B) Power spectral density in a representative postcentral electrode during a 3.5 s window centered on overt speaking or silent miming. The location of the electrode is shown on the right (red dot). (C) Distribution of spectral peaks (2–20 Hz) detected in speech responsive electrodes across SMC and SMG (N = 152, 5 subjects). Spectral decomposition and peak detection were performed using the FOOOF algorithm (*48*). (D) Left: Mean power spectrum across N=129 SMC/SMG sites that exhibited peaks within the broad theta range (5-11 Hz) showing similar overall shape across the two conditions and comparable aperiodic components. Right: Corresponding aperiodic 1/f component recovered by FOOOF. (E) Direct comparison of the central frequency, width, and height of the detected theta peaks between speaking and miming revealed no significant differences (mixed-effects analysis; p-values shown in figure; error bars represent group mean ± 95% CI). This provides direct evidence that the sensorimotor theta rhythm is not a byproduct of auditory feedback: during miming, participants moved their articulators without producing sound, yet the oscillation retained stable spectral properties comparable to overt speech. (F) Example trial showing momentary increases in theta phase coherence during both speaking (red) and silent miming (blue) across 42 speech-responsive sites in a representative subject (15 traces plotted). Voice amplitude is shown above. (G) Averaged cohereogram across N=68 pairs of speech-responsive sites from 5 subjects, demonstrating a stable increase in theta phase coherence during both speaking and silent miming. Shaded areas: 95% CI of within-electrode differences obtained from the mixed-effects analysis. (H) Spatial distribution of analyzed electrodes, color-coded by coherence during overt speech. White stars mark the reference theta sites used for phase-coherence calculation. Right-hemisphere sites are not shown.

**Figure S3.**
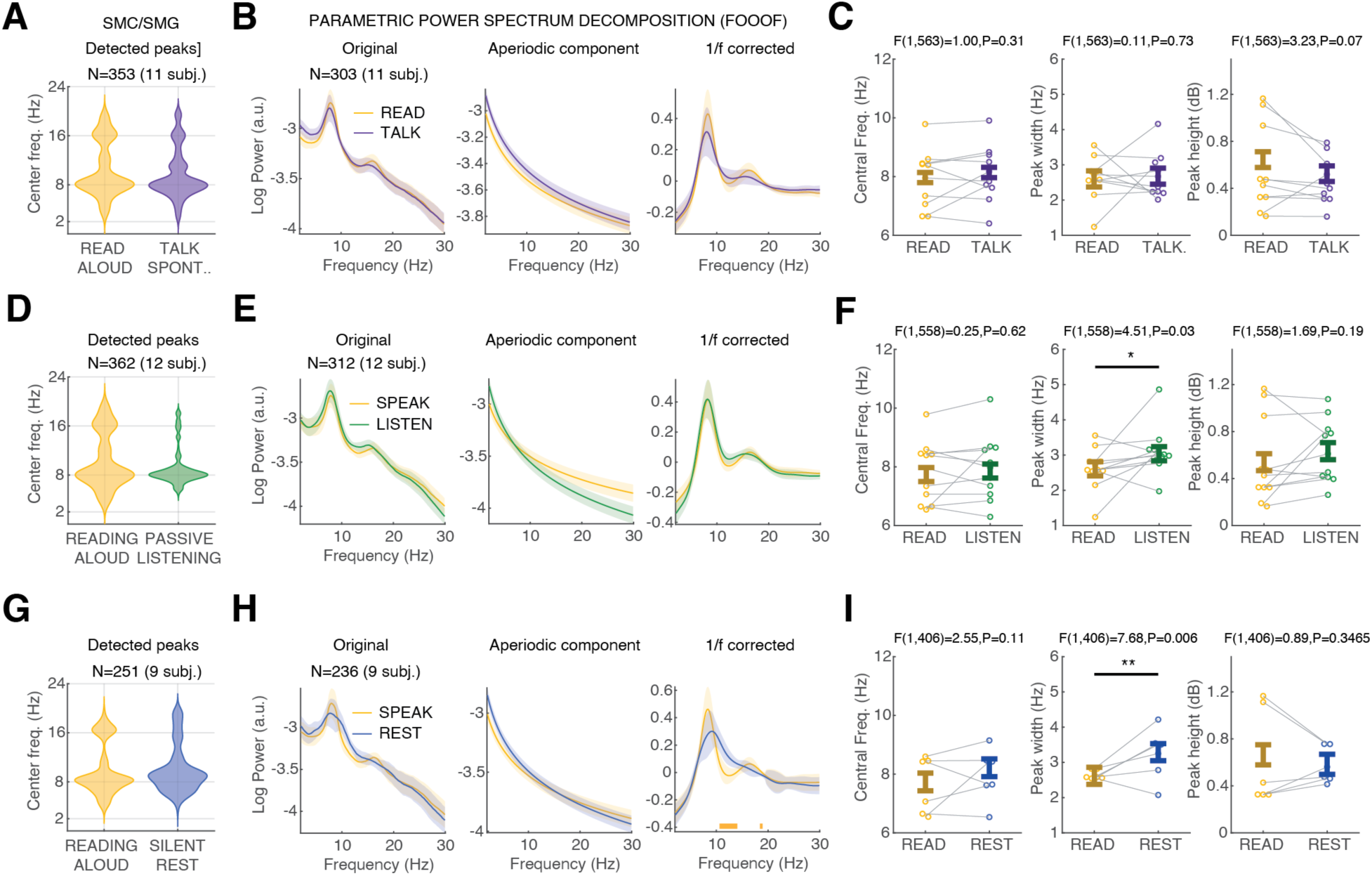
SMC theta oscillations show stability across behavioral states. All the participants (N=14) completed a control condition in which they passively listened to 225–600 sentences from the TIMIT corpus (*114*). In addition, 12 participants completed an autobiographical interview task, where they were asked to recount a personal experience or memory; from these interviews we extracted sentences spanning a broad range of natural conversational utterances. These two conditions were compared to our main sentence reading task. **(A-C)** Comparing reading aloud vs. spontaneous conversational utterances. **(D-F)** Comparing reading aloud vs. passive listening. **(G-I)** Comparing reading aloud vs. silent rest. **(A,D,G)** Distribution of spectral peaks (2–20 Hz) detected in speech responsive electrodes across SMC and SMG. Spectral decomposition and peak detection were performed using the FOOOF algorithm (*48*). **(B,E,H)** Average power spectra across electrodes exhibiting peaks within the theta range (5–11 Hz), with aperiodic and oscillatory components shown separately. Spectra were computed over –0.5 to 3.5 s windows aligned to utterance onset, revealing stable low-frequency profiles (1–30 Hz) across conditions and a prominent oscillatory peak in the theta range. ECoG time series were rescaled prior to spectral decomposition to standardize voltage values across electrodes (see Methods). **(C,F,I)** Mixed-effects analysis comparing the central frequency, width, and height of theta-range peaks. While peak frequency and magnitude remained stable, peaks were significantly broader during passive listening and silent rest (i.e., idle states of the speech motor cortex). Circles indicate subject means; error bars represent group mean ± 95% CI; p-values shown at top.

**Fig. S4.**
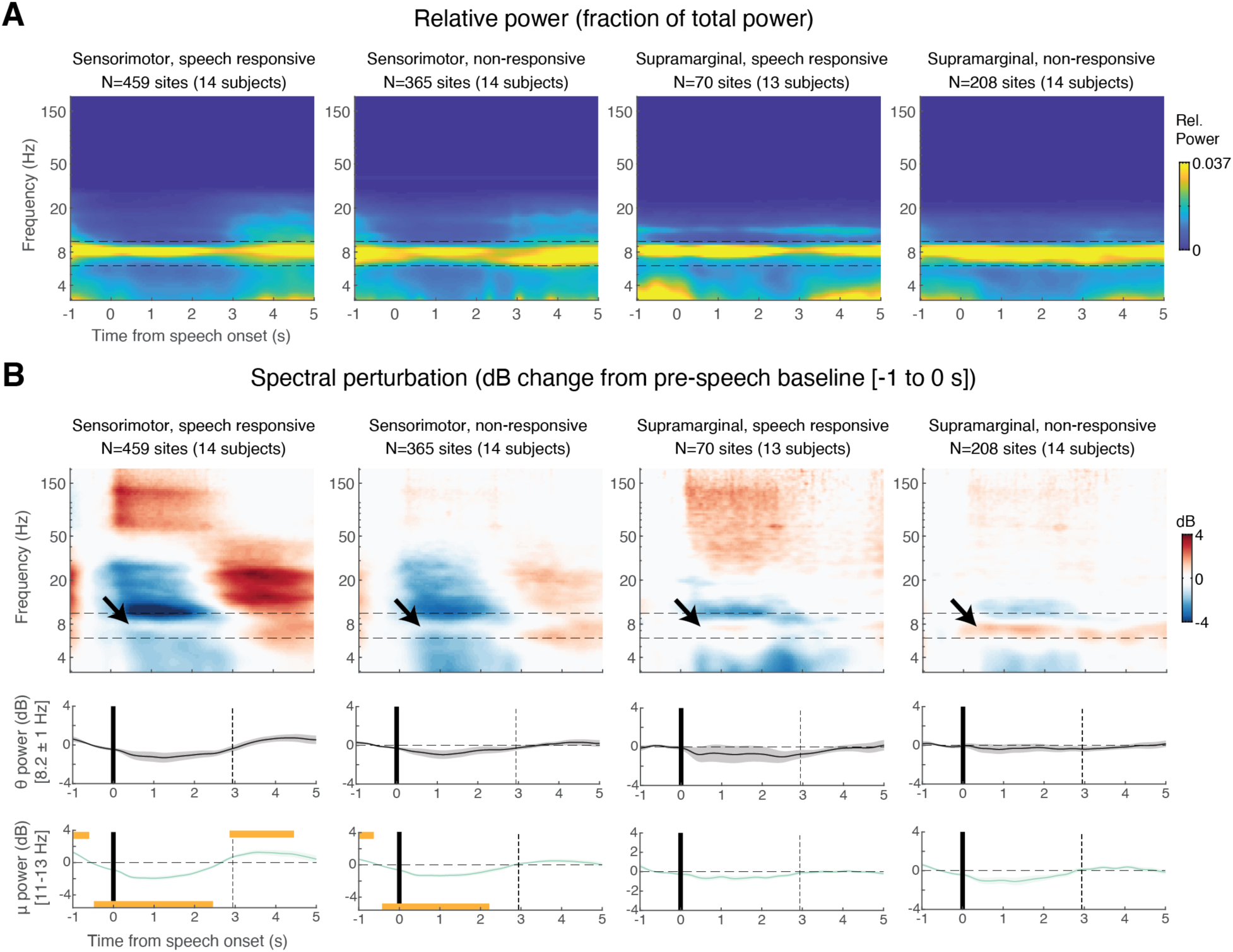
Wavelet spectrogram aligned to speech onset in SMC and SMG. Morlet wavelet spectrogram, time-locked to speech onset, showing relative power in panel **(A)** and baseline-corrected normalized power in panel **(B)**. In panel (A), colors represent the fractional power at each frequency relative to the total signal power. Notably, there is persistent oscillatory power concentrated at 8 Hz during speech, extending both before and after the utterance (mean sentence duration: 3 ± 0.4 sec). In panel (B), colors represent the dB power change relative to the immediate pre-speech baseline (−1 to 0 s). A timepoint-by-timepoint mixed-effects analysis, comparing the intercept against zero (FDR corrected), revealed no significant power changes at 8 Hz relative to speech onset (middle panels), emphasizing the persistence of the sensorimotor theta oscillation. This stability contrasts sharply with the significant post-speech power decrease in the neighboring Mu band (10–13 Hz), as indicated by the horizontal orange bars on the x-axis marking significant time points.

**Fig. S5.**
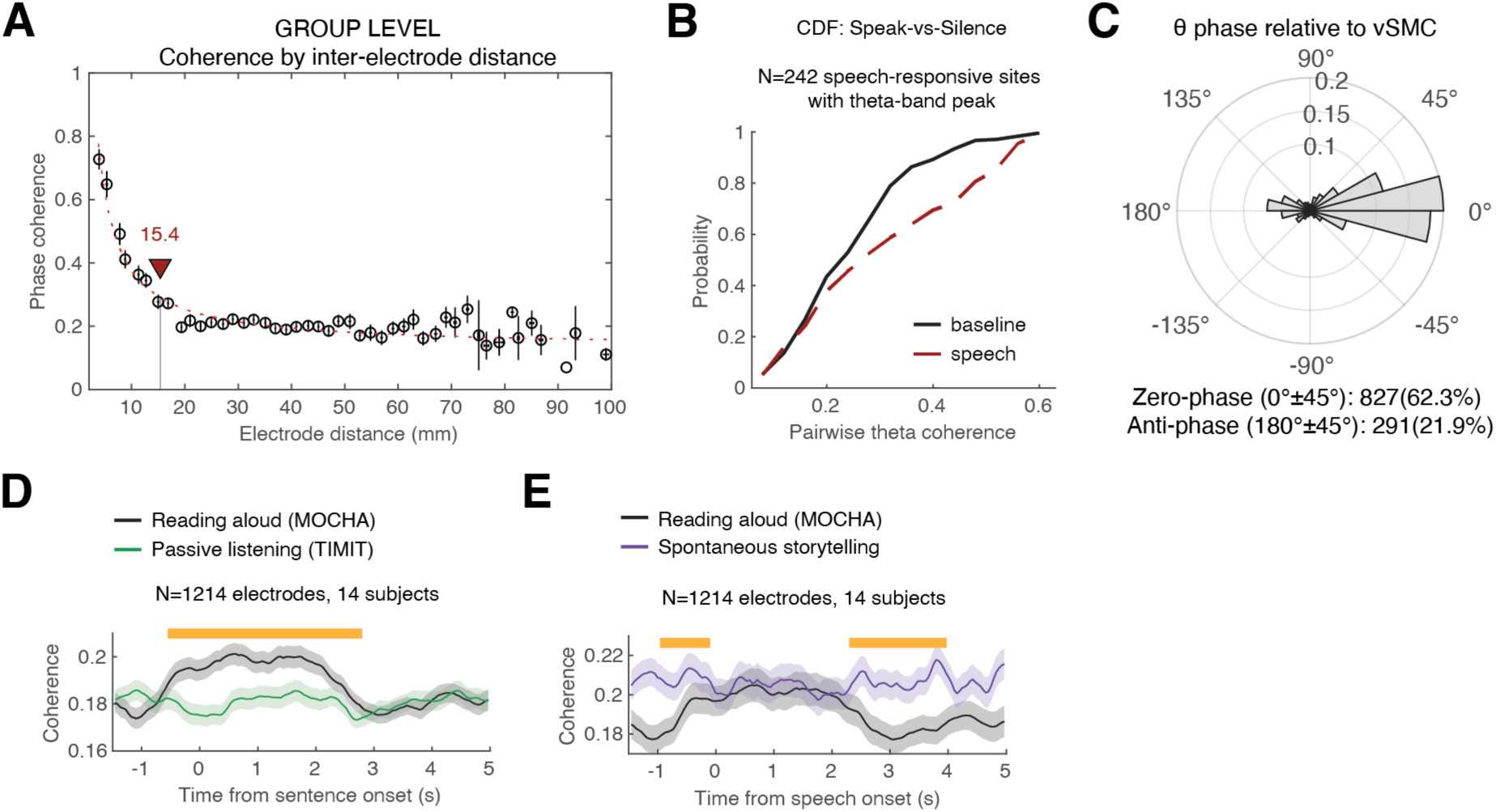
Phase coherence among speech responsive site across behavioral states. **(A)** Mean phase coherence in the theta band (6-10 Hz), computed within a time window of 0 to 4 seconds relative to speech onset, plotted as a function of inter-electrode distance (mm). Electrodes in close proximity exhibited enhanced coherence, potentially attributable to passive conductance effects. For the analysis presented in Fig. 2, we included only electrode pairs separated by at least 15 mm (where the plateau begins) to mitigate this effect. **(B)** Cumulative distribution of theta-band phase coherence during speech versus the pre-speech baseline across speech-responsive electrodes exhibiting a detectable theta peak. **(C)** Distribution of phase difference across speech responsive sites, relative to a single most ventral SMC speech-responsive electrode in each patient. Most electrodes exhibited near 0-phase coupling (0 ± 45°: 770/1328; 58%). Another substantial portion showed antiphase coupling (180 ± 45°: 277 electrodes; 21%). **(D)** Comparing theta phase coherence during speech production and passive listening revealed significantly higher coherence during production, arguing against an auditory-driven account and highlighting its motor-related origin. **(E)** Theta phase coherence during reading and spontaneous articulation showed comparable magnitudes, though spontaneous speech exhibited sustained coherence enhancement, consistent with continuous storytelling. Shaded areas: 95% CI of within-electrode differences obtained from the mixed-effects analysis. Horizontal orange markers indicate statistically significant time points.

**Fig. S6.**
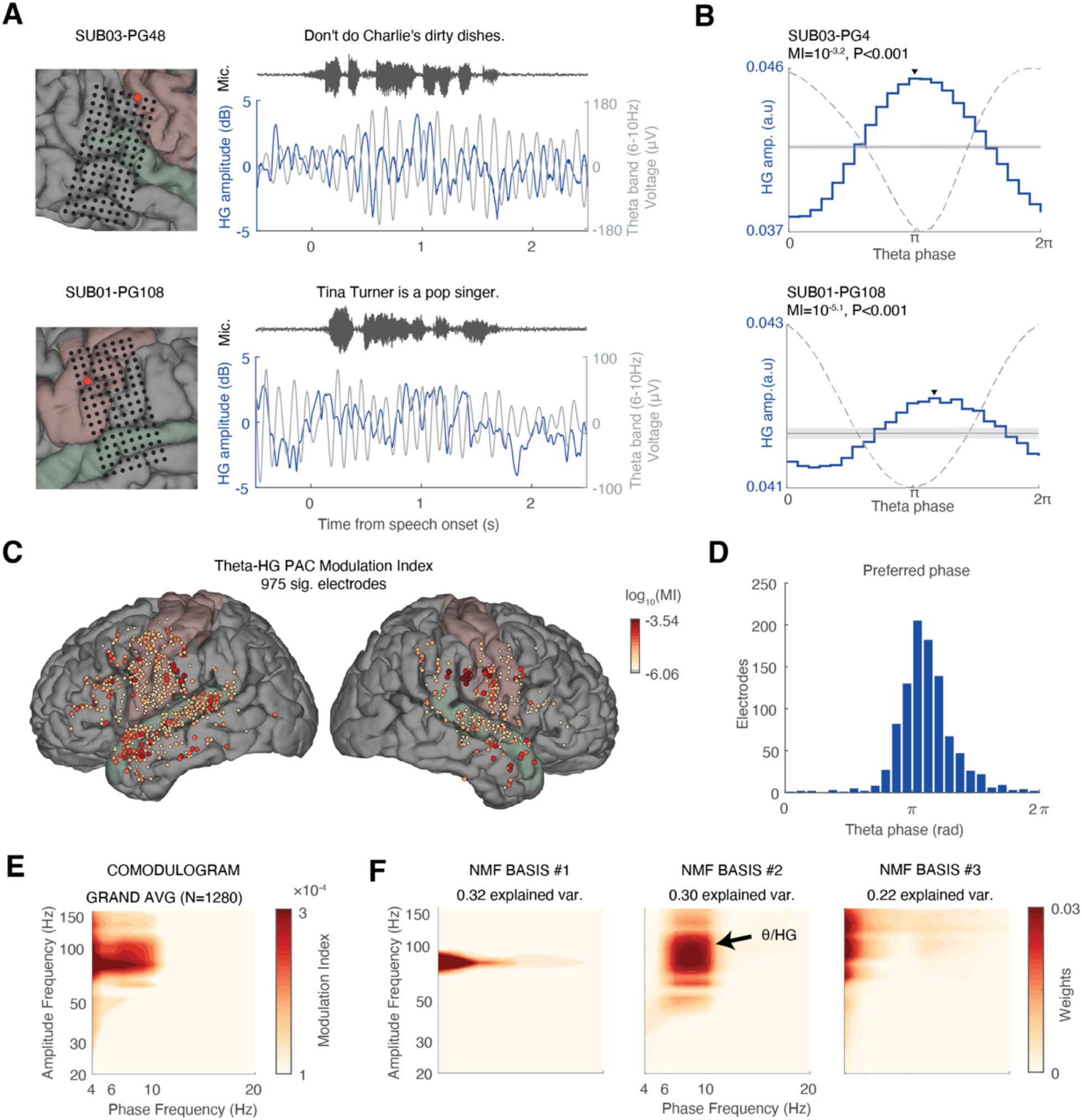
Theta-HG phase-amplitude coupling (PAC) during speech. (**A**) Example of speech-responsive vSMC electrodes from two subjects, demonstrating robust theta oscillations and phase-dependent modulation of HG power during articulation. **(B)** To quantify PAC during speech production, we used Tort’s Modulation Index (MI), as proposed by (*56*, *57*). The instantaneous theta phase was extracted via a Hilbert transform and binned into 24 phase bins spanning the range [0, 2π]. We then averaged the HG amplitude in each phase bin across all speech intervals and quantified deviations from a uniform distribution using Kullback-Leibler divergence. To statistically assess theta-HG coupling, we compared the theta-HG MI for each electrode to a surrogate distribution generated by circularly shifting the theta phase time series by a random amount 5,000 times (gray horizontal line represent the mean ± SD of the shaffled data). Significance was determined by the proportion of iterations in which the MIshuff. was equal to or greater than the empirical MI. We found that 72.1% (975/1,280) of all speech-responsive electrodes exhibited statistically significant theta-HG coupling (P<0.05, FDR corrected). **(C)** Spatial distribution of MI values across significant electrodes, highlighting the widespread nature of theta-HG PAC. **(D)** Distribution of preferred theta phases across significant electrodes, showing that HG amplitude was consistently higher near the theta trough. **(E)** Phase-amplitude comodulogram compute using the MI method, averaged across all speech-responsive electrodes (*N* = 1,280). **(F)** To decompose the grand-average comodulogram into its dominant components, we applied non-negative matrix factorization (NMF) using MATLAB’s nnmf.m function. This analysis revealed three dominant PAC components collectively accounting for over 84% of the variation across electrodes. Notably, NMF basis #2 identified a distinct PAC between theta (6–10 Hz) and HG (70–120 Hz), explaining 30% of the variance.

**Fig. S7.**
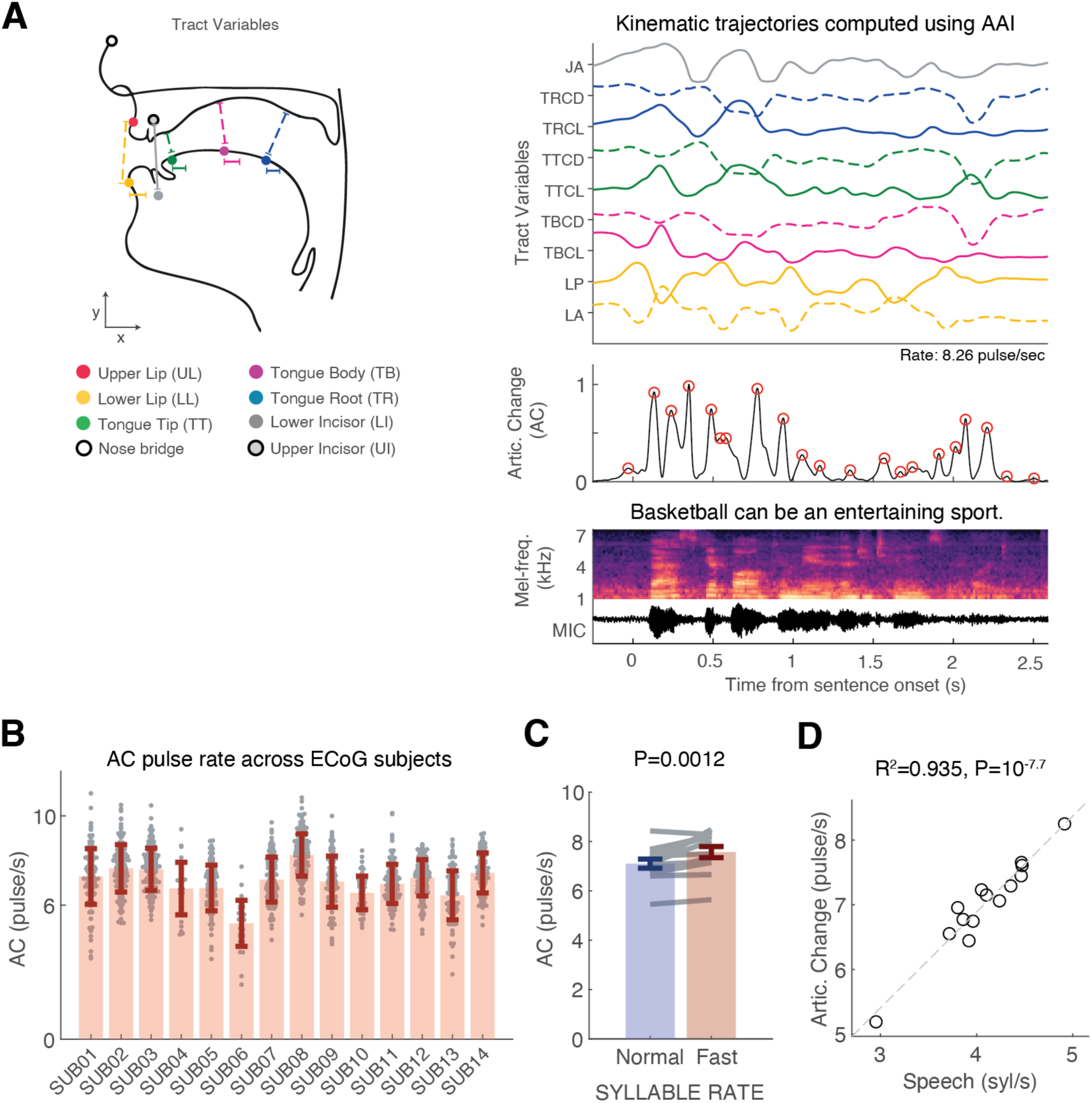
Articulatory kinematics computed through AAI in ECoG subjects. (**A**) Example of articulatory kinematics during continuous speech in a representative ECoG subject. These trajectories were inferred using AAI, based on the produced acoustics. As in actual EMA measurements, articulatory movements exhibited semi-rhythmic modulation, characterized by abrupt, pulse-like changes in the sum of squared velocities across articulators (AC). **(B)** AC pulse rates across 14 ECoG subjects. Each gray dot represents a single sentence; error bars indicate mean ± SD. **(C)** Similar to the pattern observed in the EMA dataset, AC pulse rates in ECoG subjects were higher during faster-spoken sentences (P<0.01, Wilcoxon singed-rank test), yet remained centered within the theta range. For normally spoken sentences (10th–90th syllable rate percentiles): mean syllable rate = 4.2 syl/s, AC pulse rate = 7.1 ± 0.18 Hz. For the fastest sentences (>90th percentile): mean syllable rate = 5.5 syl/s, AC pulse rate = 7.6 ± 0.23 Hz). **(D)** Across subjects, AC pulse rate was strongly correlated with the mean syllable rate, highlighting the tight correspondence between these motor events and the resulting speech sounds (R^2^=0.93, P < 10⁻⁷).

**Fig. S8.**
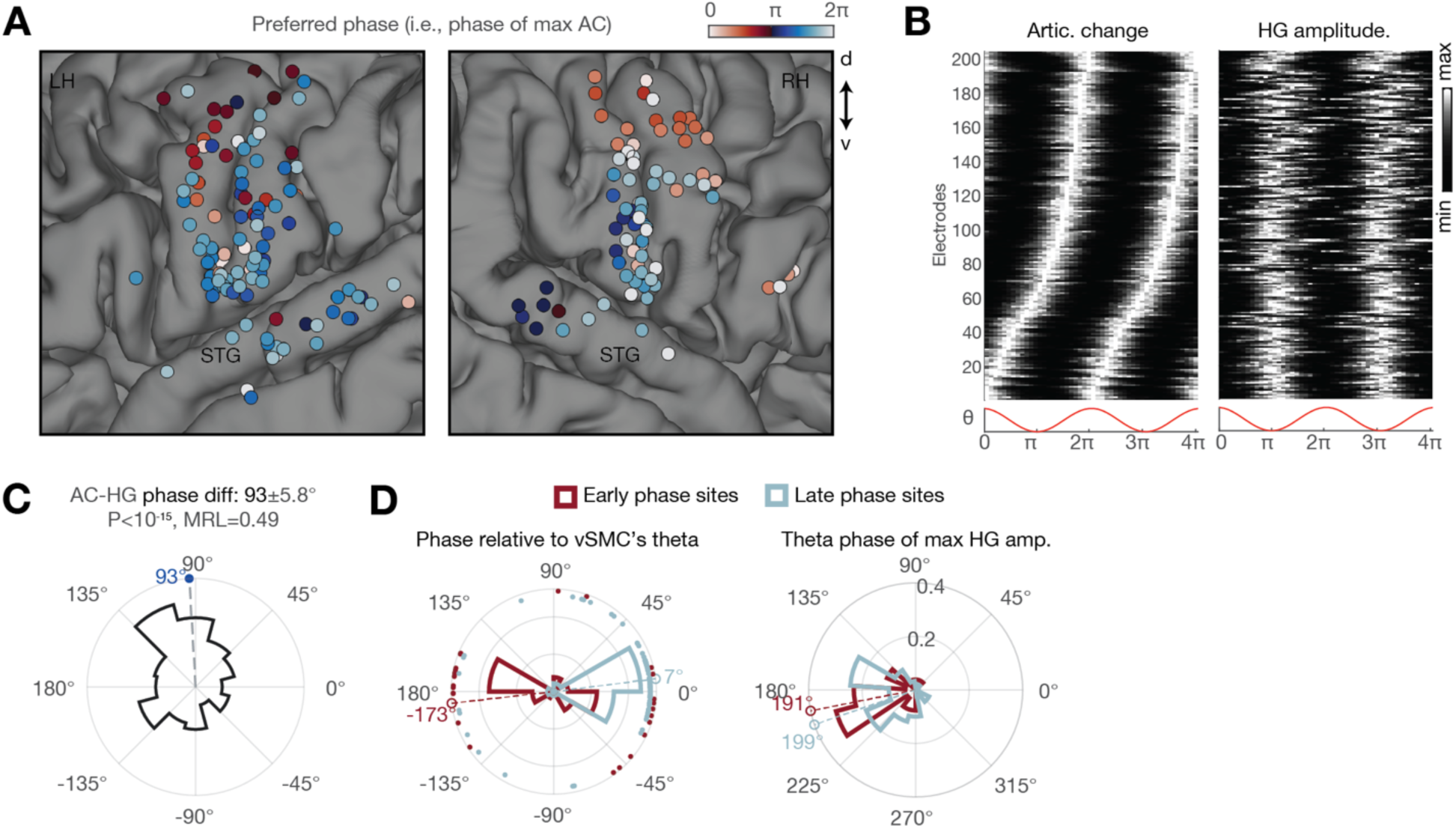
Consistent phase difference along the SMC dorsoventral axis. **(A)** Anatomical distribution of the preferred theta phase (i.e., phase at which AC is maximal) across electrodes with significant coupling (P<0.05, FDR corrected). Notice a cluster of electrodes in the dorsal SMC manifesting coupling to an earlier phase of the theta cycle. **(B)** The electrodes plotted in (A), sorted by their preferred theta phase (phase of maximal AC). Right: the same electrodes, same sorting, but for HG amplitude. Despite variability in the preferred phase for AC, the electrodes exhibit a consistent HG amplitude increase around the theta trough. **(C)** Distribution of the within-electrode difference between theta phase of maximal HG and maximal AC (angular difference: 93° ± 5.8°, mean resultant length **|**R**|**=0.49, P<10-15 Rayleigh’s test). **(D)** To assess whether the theta rhythm itself was phase-shifted at sites showing early coupling in (B), we calculated the phase difference between each electrode’s theta signal and that of the most ventral SMC electrode in each patient, which served as a consistent reference site across subjects (both anatomically and functionally). The analysis revealed that most sites where AC was coupled to an earlier phase of the theta cycle (i.e., before the trough, from ½π to π) exhibited nearly antiphase theta rhythm (−173±15.1°; early-vs-late electrodes: P<0.001) compared to the “late-coupling” more ventral sites (i.e., after the trough, from ³⁄₂π to 2π). Importantly, within each site, the HG amplitude was still consistently coupled to the trough (right panel; early-vs-late electrodes: P=0.52 N.S).

**Fig. S9.**
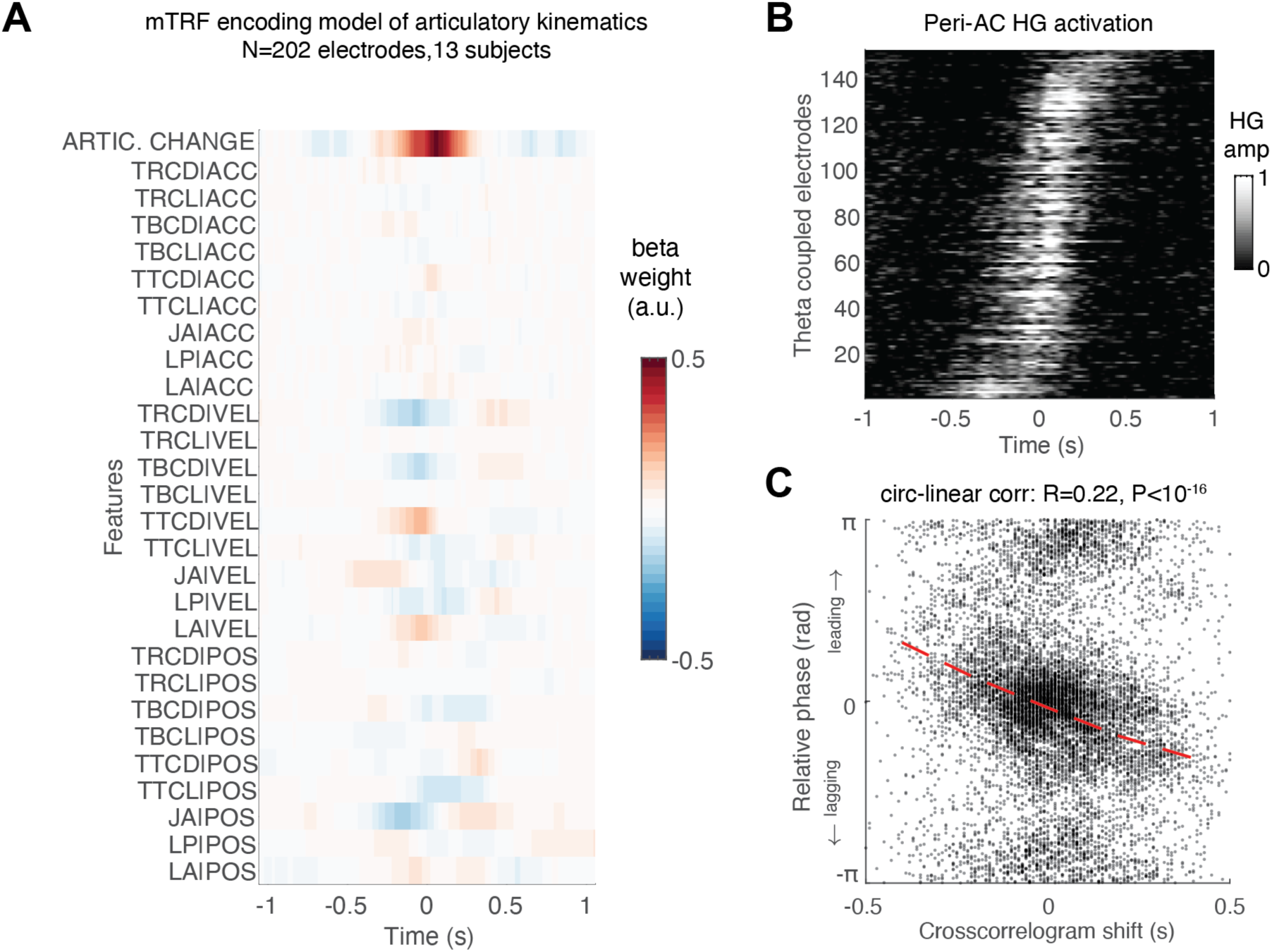
Mean encoding model beta weights and correlation between peri-AC responses and theta phase. **(A)** Beta weights obtained from the mTRF articulatory encoding model, fitted to electrodes exhibiting significant theta-movement coupling (P < 0.05, FDR corrected; N = 202 electrodes across 13 subjects). Note the prominent beta weights corresponding to the Articulatory Change feature. This feature stands out among other kinematic features, explaining a significant amount of unique variance in more than 60% of the electrodes (P < 0.05, compared to a null distribution generated by circularly shifting the AC feature by a random amount 200 times and refitting the model). Across subjects, this unique variance was significantly greater than zero (proportion of variance: 1 ± 0.09%, P < 0.0004, Wilcoxon signed-rank test, N = 14 subjects). **(B)** Across the electrodes exhibiting significant unique variance explained by AC, we observed a progression of response latencies, with some electrodes activating slightly before and others slightly after the AC signal. **(C)** Correlation between the time differences among electrodes— determined by the peaks of the cross-correlograms of peri-AC responses—and their relative theta phase differences (computed in reference to the most ventral SMC site in each subject). We found significant circular-linear correlation across electrode pairs (R=0.22, P<10^-16^). Dashed red line, running circular mean.

**Fig. S10.**
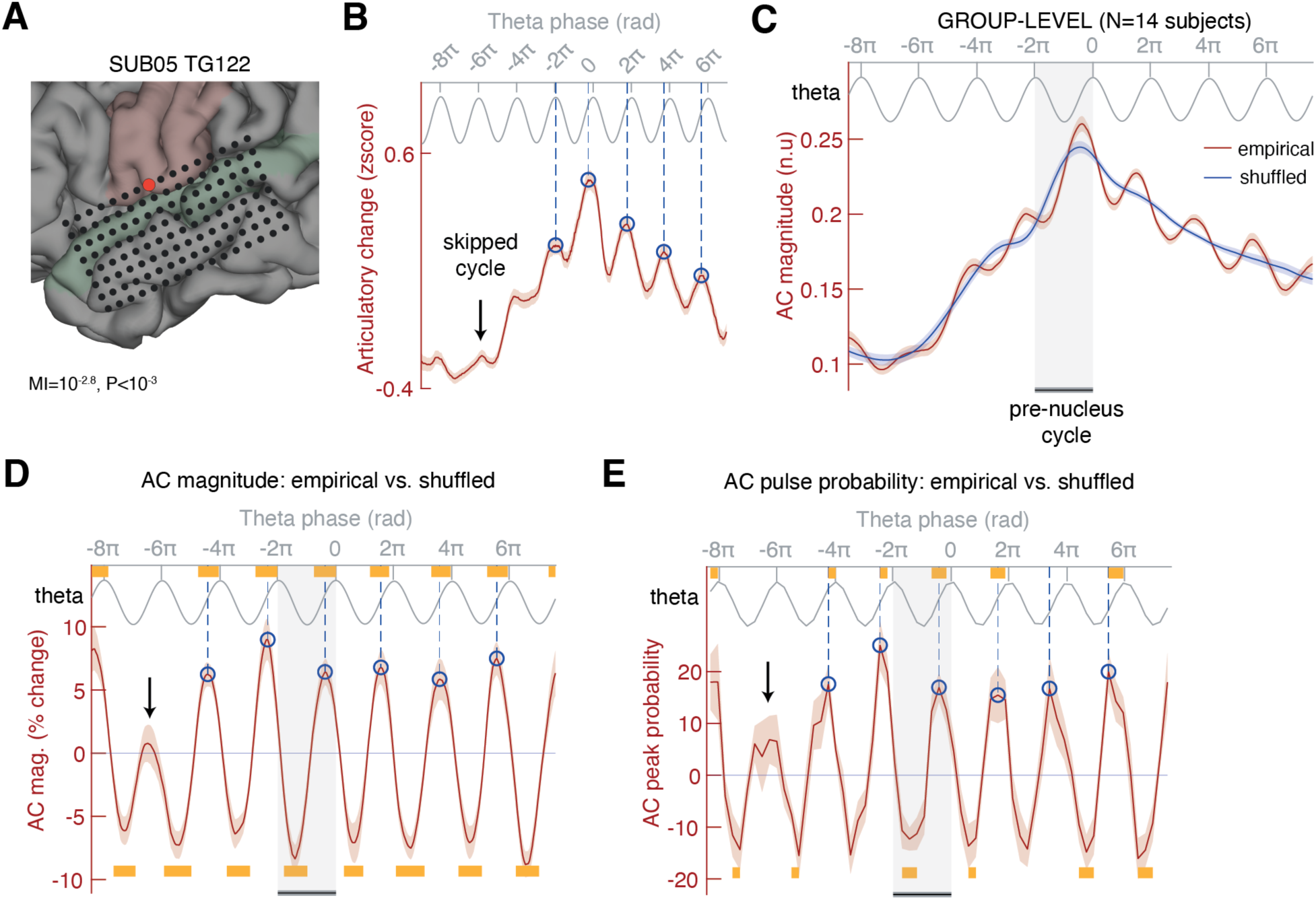
Theta oscillation coordinates successive vocal-tract movements during syllable production. (A–B) Same analysis as described in Fig. 6, shown for an electrode from another subject. **(C–E)** Group-level analysis (one electrode per subject, selected for strongest theta–movement coupling) shows that successive AC pulses during syllable production are precisely timed by the ongoing theta rhythm, emerging shortly after the trough and before the subsequent peak. Panel C shows the grand-average AC profile alongside a surrogate profile generated by circularly shifting theta phase in each trial by a random amount (see Methods). Panel D presents AC magnitude expressed as percent change from shuffled data. A cluster-based permutation test revealed significant AC modulation by theta phase (yellow bars indicate statistical significance). The dashed blue line marks the average phase of individual AC peaks, highlighting their consistent alignment with a specific theta phase. Panel E shows a theta phase histogram of AC peaks, displaying the empirical probability distribution normalized by the surrogate distribution. Yellow bars indicate theta phases where the probability of AC pulse was significantly increased/decreased. Shaded areas represent ±SE across subjects. The persistence of this coupling—even after brief pauses in speech (black arrows) during which an entire theta cycle was skipped—further supports the idea that the AC pulses ride on top of the sensorimotor oscillation; that is, the oscillation governs the timing of articulatory movements rather than being driven by them.

**Fig. S11.**
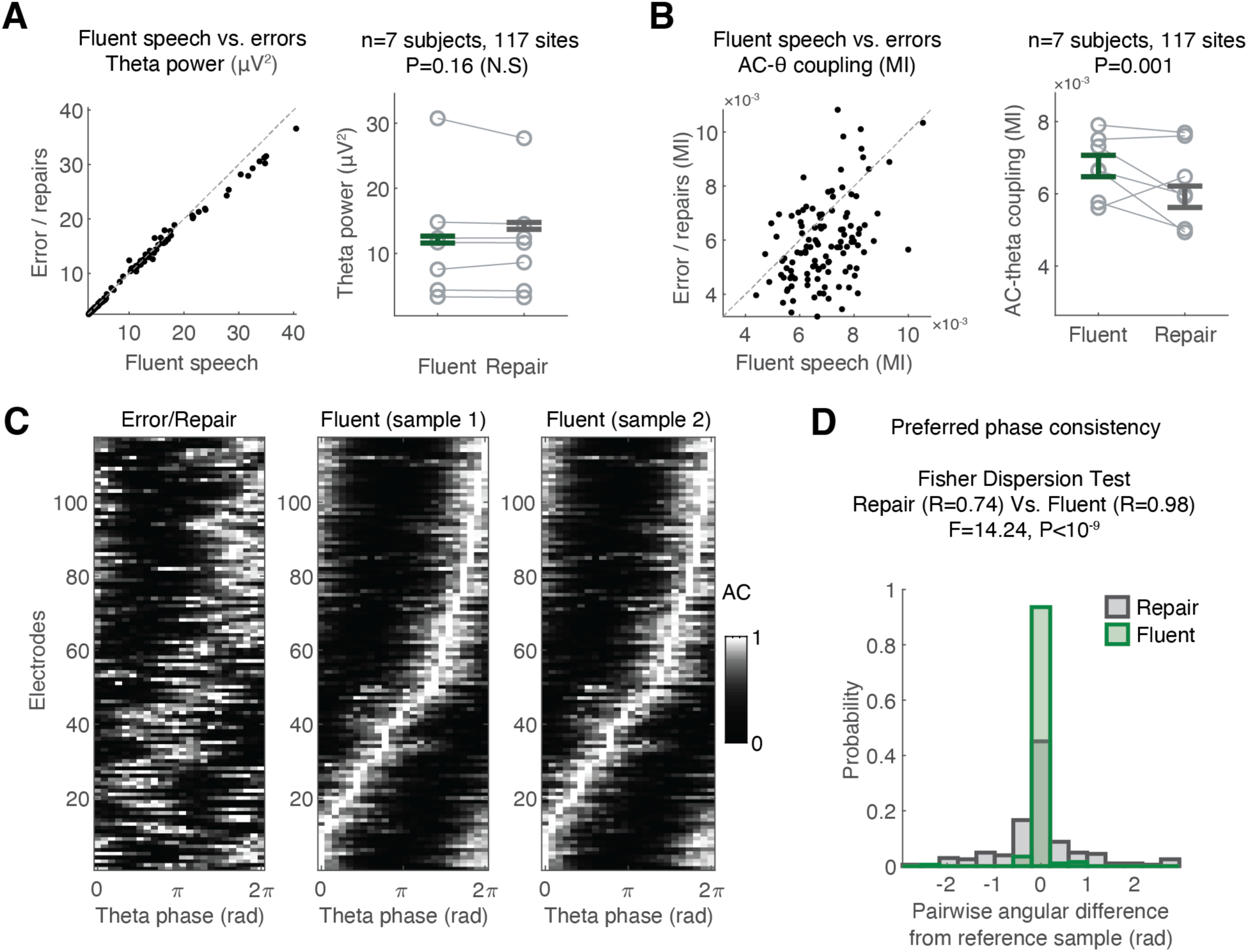
Disruption of Theta-Movement Coupling During Speech Errors. We compared theta power and theta-movement coupling strength between fluent speech and speech errors/repairs. Subjects with fewer than 10 errors were excluded. To create a comparable fluent speech group with the same number of sentences, we randomly sampled trials while matching syllable rate and sentence duration to the error group. This was repeated 200 times, and median values were used for comparison. **(A)** Theta Power: Scatter plot (right) compares theta power between fluent and error trials across electrodes with significant theta-movement coupling. Mixed effects analysis (left) showed no significant difference (P = 0.16, mixed effects analysis, 117 sites, 7 subjects). **(B)** Theta-Movement Coupling (MI): Scatter plot (right) compares theta-movement coupling between fluent and error trials. Mixed effects analysis (left) showed a significant difference (P=0.001, mixed effects analysis, 117 theta-coupled sites across 7 subjects). **(C)** Preferred phase across electrodes during speech errors/repairs versus fluent sentences (using odd and even samples). Electrodes are sorted based on the even samples (sample 2). The preferred phase remained highly consistent across the two fluent speech samples, whereas speech errors/repairs exhibited a more variable distribution. **(D)** Comparing the (within-electrode-) angular difference between the preferred phase found in odd vs. even samples of fluent speech to the difference found between errors vs. fluent speech revealed greater phase-of-movement variability during trials with errors/repairs (resultant vector length: errors/repairs = 0.73, fluent = 0.99; P < 10⁻⁶, Fisher dispersion test).

**Fig. S12.**
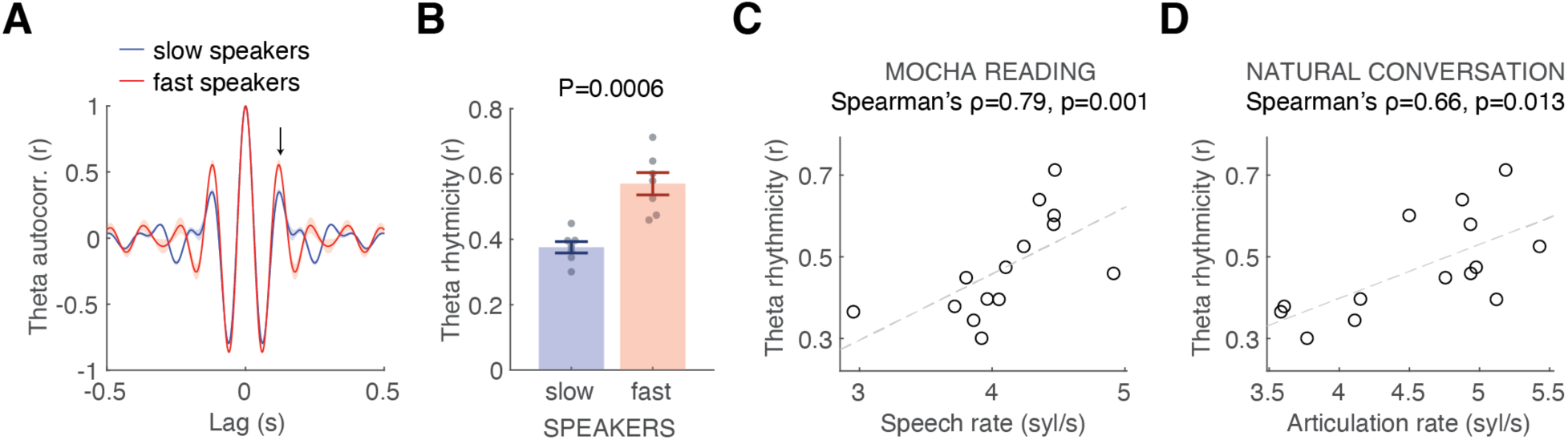
Faster speech is associated with stronger theta rhythmicity. We analyzed the autocorrelation function (ACF) of the theta-band signal from the top five electrodes showing robust theta–movement coupling. **(A)** We divided the 14 subjects into fast and slow speakers using a median split based on their mean syllable rate. Fast speakers exhibited more pronounced rhythmicity, reflected by a higher secondary peak in the ACF (black arrow). **(B)** Theta rhythmicity was significantly stronger in fast speakers (*P* = 0.0006, rank-sum test). **(C-D)** We observed a significant correlation between median syllable rate and theta rhythmicity both during the MOCHA reading task and during natural conversation. For the latter, we analyzed spontaneous utterances (mean length: 11 ± 3.1 words per utterance) and computed the median articulation rate across utterances (syllables per second, excluding pauses). One subject lacked natural conversation data; for this individual, we used held-out sentence reading data that had not been included in the theta analysis.

